# Loss of the lysosomal protein CLN3 modifies the lipid content of the nuclear envelope leading to DNA damage and activation of YAP1 pro-apoptotic signaling

**DOI:** 10.1101/2024.05.31.596474

**Authors:** Neuza Domingues, Alessia Calcagni’, Joana Pires, Sofia Roque Freire, Niculin Joachim Herz, Tuong Huynh, Katarzyna Wieciorek, Maria João Moreno, Tiago Fleming Outeiro, Henrique Girão, Ira Milosevic, Andrea Ballabio, Nuno Raimundo

**Affiliations:** Multidisciplinary Institute of Ageing, University of Coimbra, Coimbra, Portugal; Telethon Institute of Genetics and Medicine (TIGEM), Naples, Italy; Department of Translational Medical Sciences, Federico II University, Naples, Italy; Department of Molecular and Human Genetics, Baylor College of Medicine, Houston, TX, USA; Jan and Dan Duncan Neurological Research Institute, Texas Children’s Hospital, Houston, TX, USA; University Medical Center Göttingen, Department of Experimental Neurodegeneration, Center for Biostructural Imaging of Neurodegeneration, Göttingen, Germany; CQC-Biological Chemistry Group, Chemistry Department FCTUC, Coimbra, Portugal; Translational and Clinical Research Institute, Newcastle University, Newcastle upon Tyne, UK; Max Planck Institute for Multidisciplinary Sciences, Göttingen, Germany; Deutsches Zentrum für Neurodegenerative Erkrankungen (DZNE), Göttingen, Germany; Coimbra Institute for Clinical and Biomedical Research (iCBR), Centre for Innovative Biomedicine and Biotechnology, Academic and Clinical Center of Coimbra, Faculty of Medicine, University of Coimbra, Portugal; Centre for Human Genetics, Nuffield Department of Medicine, University of Oxford, UK; SSM School for Advanced Studies, Federico II University, Naples, Italy; Department of Cellular and Molecular Physiology, Penn State College of Medicine, Hershey, PA, USA; Penn State Cancer Institute, Hershey, PA, USA

## Abstract

Batten disease is characterized by early-onset blindness, juvenile dementia and death during the second decade of life. The most common genetic causes are mutations in the *CLN3* gene encoding a lysosomal protein. There are currently no therapies targeting the progression of the disease, mostly due to the lack of knowledge about the disease mechanisms.

To gain insight into the impact of CLN3 loss on cellular signaling and organelle function, we generated CLN3 knock-out cells in a human cell line (CLN3-KO), and performed RNA sequencing to obtain the cellular transcriptome. Following a multi-dimensional transcriptome analysis, we identified the transcriptional regulator YAP1 as a major driver of the transcriptional changes observed in CLN3-KO cells.

We further observed that YAP1 pro-apoptotic signaling is hyperactive as a consequence of CLN3 functional loss in retinal pigment epithelia cells, and in the hippocampus and thalamus of CLN3^exΔ7/8^ mice, an established model of Batten disease. Loss of CLN3 activates YAP1 by a cascade of events that starts with the inability of releasing glycerophosphodiesthers from CLN3-KO lysosomes, which leads to perturbations in the lipid content of the nuclear envelope and nuclear dysmorphism. This results in increased number of DNA lesions, activating the kinase c-Abl, which phosphorylates YAP1, stimulating its pro-apoptotic signaling.

Altogether, our results highlight a novel organelle crosstalk paradigm in which lysosomal metabolites regulate nuclear envelope content, nuclear shape and DNA homeostasis. This novel molecular mechanism underlying the loss of CLN3 in mammalian cells and tissues may open new c-Abl-centric therapeutic strategies to target Batten disease.

## INTRODUCTION

Neuronal ceroid lipofuscinoses (NCLs), also referred as Batten disease, are the most prevalent neurodegenerative lysosomal storage diseases (LSD) (Platt et al., 2018). The pathology of NCLs exhibits devastating manifestations such as blindness, seizures or epilepsy, dementia, loss of cognitive and motor skills, ultimately leading to death around 20 years of age (Simonati & Williams, 2022). There are currently no therapies targeting the disease progression, mostly due to the lack of knowledge about the disease mechanisms. The NCLs are characterized by mutations in 13 genes encoding different CLN) proteins, e.g., CLN2, CLN3, CLN5. The most common mutations are in the gene encoding CLN3 (Mole & Cotman, 2015), particularly a 1.02 kb deletion which removes exons 7 and 8 (CLN3^Δex7/8^). This mutation results in an open-reading frame disruption that leads to either mRNA decay, or translation of a truncated protein at the C-terminus (Centa et al., 2023; Miller et al., 2013).

CLN3 is an ubiquitously-expressed multi-pass transmembrane protein initially reported to be predominantly localized at the lysosomal membrane (Mirza et al., 2019) and, more recently, at the Golgi apparatus (Calcagni’ et al., 2023). Additionally, CLN3 was shown to regulate the trafficking of cation-independent mannose-6-phosphate receptor (CI-M6PR), revealing an important function in the secretion of lysosomal enzymes, and in autophagic-lysosomal reformation (Calcagni’ et al., 2023). CLN3 was also shown to be involved in the export of glycerophosphodiesters from the lysosomes (Laqtom et al., 2022). Glycerophosphodiesters originate from the lysosomal degradation of phospholipids, which are first metabolized into lysophospholipids, and then further converted into other glycerophosphodiesters. Interestingly, brain tissue from mice lacking CLN3 revealed lysosomal accumulation of phospholipid catabolism products, including the degradation intermediate lysophosphatidylglycerol (Laqtom et al., 2022). Despite this implication of CLN3-loss of function in lysosomal reformulation and cellular lipid homeostasis, its effects on the dynamics of other organelles and the subsequent impact on cellular signaling remain unclear.

Organelle crosstalk is vital for maintaining cellular homeostasis, and its disruptions interfere with different signaling pathways that lead to cellular dysfunction and to pathological conditions. Lysosomes communicate with various organelles, including mitochondria (Deus et al., 2019; Fernandez-Mosquera et al., 2019; Yambire, Mosquera, et al., 2019), endoplasmic reticulum (ER) (Höglinger et al., 2019) and peroxisomes (Chu et al., 2015). Lysosomes communicate with the nucleus through signaling pathways (e.g., lysosomal biogenesis induced by the microphtalmia transcription factors such as TFEB (Sardiello et al., 2009; Settembre et al., 2011)). In *C. elegans*, upon induction of lysosomal lipolysis, fatty acid-binding protein homolog-8 was shown to translocate from the lysosome to the nucleus (Folick et al., 2015), delivering oleoylethanolamide to the nuclear hormone receptor complex NHR-49-NHR-80, which induces the expression of mitochondrial metabolic genes (Ramachandran et al., 2019). No membrane contact sites have been described between lysosomes and nucleus in mammalian cells. Despite the enormous potential of lipids as molecular candidates to mediate lysosome-nucleus communication, this process remains poorly understood.

Here, we demonstrate that the loss of CLN3 function depletes the nuclear envelope of key lipid species, causing nuclear dysmorphism and DNA damage, which triggers a pro-apoptotic response mediated by the transcription factor YAP1. These findings were observed in different human cell lines and in the brain of a mouse model of Batten disease.

YAP1 is an important regulator of cell proliferation and death, and is considered a canonical effector of the Hippo pathway (Zhao et al., 2010). In addition to this canonical regulation of YAP1 activity through the Hippo pathway, other kinases phosphorylate YAP1, regulating its localization, interaction with co-activators and transcriptional activity. Accordingly, in response to DNA damage, tyrosine kinase c-Abl relocates into the nucleus and phosphorylates YAP1 on Tyr357 (Levy et al., 2008), which stimulates YAP1 interaction with tumor suppressor p73 and a pro-apoptotic gene expression program (Lapi et al., 2008; Strano et al., 2005). In addition, in the context of LSDs, YAP1 was also found to interact with TFEB, contributing to accumulation of autophagosomes (Ikeda et al., 2021). The activation of a pro-apoptotic signaling pathway caused by impaired lipid trafficking between lysosomes and nuclear envelope opens a new paradigm for inter-organelle communication and cellular signaling.

## RESULTS

### Loss of CLN3 in human cells triggers YAP signaling

To determine the signaling consequences of loss of CLN3, we generated CLN3 knockout in human embryonic kidney (HEK) 293 cells (henceforth CLN3-KO) with CRISPR-mediated deletion of 10 nucleotides in the CLN3 gene (**Supplementary Figures S1A**), which causes premature termination (**Supplementary Figure S1B**). The protein levels of CLN3 were strongly reduced the CLN3-KO cells (**Supplementary Figure S1C**). First, we confirmed the expected lysosomal phenotypes of CLN3 loss-of-function (Calcagni’ et al., 2023) in this cell line by evaluating lysosomal morphology. Upon staining the cells with anti-Lamp1 antibody to mark the lysosomal membranes, we observed that size of the Lamp1-positive particles was robustly increased in CLN3-KO (**Supplementary Figure S1D**), while the total number of lysosomes was similar in CLN3-KO and wild-type (WT, **Supplementary Figure S1E**). In agreement with this result, expression of Lamp1 protein in CLN3-KO cells was unchanged (**Supplementary Figure S1F**). To confirm that the increase in lysosomal size was a specific consequence of CLN3 loss rather than an adaptive mechanism, we re-expressed CLN3 (CLN3-GFP) in the CLN3-KO cells, where it localized to the lysosomes as expected (**Supplementary Figure S1G**). Expression of CLN3-GFP was sufficient to rescue the ‘swollen’ phenotype of CLN3-KO lysosomes (**Supplementary Figure 1H**). We next investigated lysosomal function in CLN3-KO cells by assessing lysosomal proteolytic activity and pH. We loaded the cells with the pH-insensitive AlexaFluor647-dextran and the pH-sensitive FITC-labelled dextran, or with Magic Red, a readout of cathepsin B activity. Of note, bafilomycinA1 treatment was used as positive control for non-acidic lysosomes. We observed that the CLN3-KO cells had lower cathepsin B activity and higher pH than the WT cells (**Supplementary Figure S1I**). Altogether, these results suggest that, while the lysosomal mass does not seem to be altered in CLN3-KO cells, their lysosomes are dysfunctional.

To determine the key consequences of CLN3-KO in an unbiased manner, we performed bulk RNA sequencing (**Figure 1A**). We identified 2288 differentially expressed genes (DEGs), with 909 up-regulated and 1379 down-regulated in CLN3-KO (**Supplementary Figure 1B** and **Supplementary Table S1**). We then analyzed the pathways predominantly enriched in the DEG list, and observed that the top pathways were related to cell proliferation (cell cycle, mTORC1 signaling), retinoic acid signaling, and DNA damage (**Figure 1B**). Next, we determined which transcription factors (TFs) were predicted to mediate the changes observed in the transcriptome. The only two transcription factors predicted to be hyperactive in CLN3-KO cells are YAP1 and SMAD1 (**Figure 1C**). The enrichment of YAP1 targets in the CLN3-KO DEG list was much higher than for SMAD1, hence we focused on YAP1 for mechanistic follow-up.

**Figure 1.**
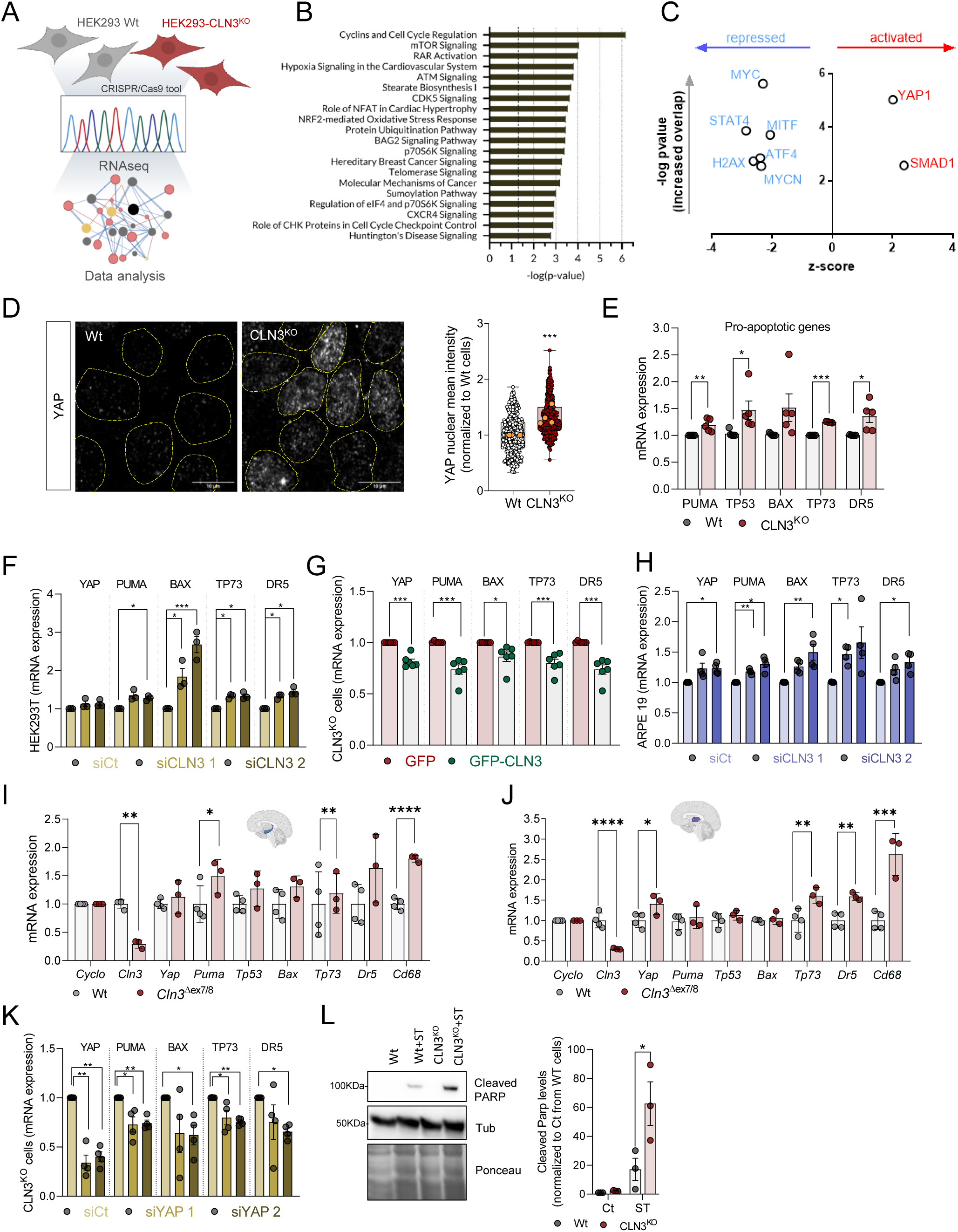
YAP signalling pathway is activated in CLN3^KO^ cells and drives a pro-apoptotic phenotype. (**A**) Schematic representation of the experimental design. (**B**) Pathway enrichment analysis of the differentially expressed genes (DEG). (**C**) Transcription factors predicted from the DEG list (**B**) represented on a volcano plot of z-score (y-axis) versus p-value of target enrichment on the DEG (x-axis) (**D**) Confocal fluorescence images of Wt and CLN3^KO^ cells immunostained for YAP protein. Nuclei are outlined by the yellow dashed lines using Hoechst staining. Scale bar 10µm. On the right, quantification of the mean intensity of YAP at the nucleus of at least 100 cells per each conditions. The yellow dots represent the mean of each independent experiment. (**E-H**) mRNA levels of YAP target genes involved in apoptosis in HEK293T WT and CLN3^KO^ stable cell lines (**E**), in WT cells with transient CLN3-depletion using two different siRNA (**F**), CLN3^KO^ cell line transfected with GFP or GFP-CLN3 and Wt cell line using two different siRNA (**G**). (**H**) mRNA levels of YAP target genes involved in apoptosis in ARPE19 cell line transfected with control (siCt) or two different siRNA directed to CLN3 transcript. (**I-J**) mRNA levels of YAP-target pro-apoptotic genes in hippocampus (**I**) and thalamus (**J**) tissue collected from 12 months old control (Wt) and *Cln3*^Δex7/8^ animals. (**K**) mRNA levels of YAP target genes involved in apoptosis in HEK293T CLN3^KO^ cell line transfected with control (siCt) or two different siRNA directed to YAP transcript. (**L**) Representative immunoblot of cleaved PARP and quantification of HEK293T (Wt and CLN3^KO^) cells with or without staurosporine (ST) treatment (4 hours). All the results are mean±SEM of at least three independent experiments. Statistical analysis with one-way ANOVA followed by Sidak’s multiple comparisons test (**E**), Tukey’s post-test (**F, H, K**), and Student’s *t*-tests when comparing two experimental groups. * p<0.05; ** p<0.01; ***p<0.001.

### YAP1 pro-apoptotic signaling is increased in CLN3-KO cells

Considering that YAP1 intracellular location is tightly regulated as it shuffles between nucleus and cytoplasm, with nuclear location being necessary for its activity, we first determined the intracellular distribution of YAP1. We observed a robust increase in the nuclear localization of YAP1 in CLN3-KO cells (**Figure 1D**), consistent with increased YAP1 activity in these cells.

YAP1 signaling affects predominantly the balance between cell proliferation and cell death (Piccolo et al., 2023). Our transcriptome analysis revealed several pathways related with DNA damage, a context that promotes YAP1 pro-apoptotic signaling (Levy et al., 2008). Notably, a key feature of Batten disease is the loss of neurons and retinal cells by apoptosis (Piccolo et al., 2023). Therefore, we focused on YAP1 pro-apoptotic signaling first. We compared the expression levels of several YAP1 transcriptional targets that are involved in pro-apoptotic signaling (PUMA, TP53, TP73, BAX, DR5), and observed that these transcripts were present at higher levels in CLN3-KO cells (**Figure 1E**), supporting increased YAP1 transcriptional activity. To confirm that the YAP1-associated pro-apoptotic signaling is a direct consequence of CLN3 loss and not a long-term adaptation, we performed a transient knock-down of CLN3 in HEK cells, using two siRNA constructs to control for off-target effects. This resulted in a decrease of CLN3 transcripts (**Supplementary Figure 2A**) and CLN3 protein levels (**Supplementary Figure 2B**), as well as ‘swollen’ lysosomes (**Supplementary Figure 2C**), indicating that the silencing of CLN3 had effectively perturbed lysosomes. We observed an increase in the transcript levels of pro-apoptotic YAP1 targets in CLN3-silenced HEK cells (**Figure 1F**), suggesting that YAP1-driven pro-apoptotic signaling is a direct consequence of CLN3 loss. To further test this premise, we re-expressed CLN3 in CLN3-KO cells, and observed a decrease in the expression levels of YAP1 transcriptional targets (**Figure 1G**).

We then sought to test if loss of CLN3 also triggered pro-apoptotic signaling in a cell type directly involved in the disease pathology. To that end, we used ARPE19 cells, a human model of retinal pigmental epithelium which degenerates in Batten disease (Calcagni’ et al., 2023; Puranam et al., 1997). In CLN3-silenced ARPE19 cells (**Supplementary Figure S2D-F**), we observed an increase in the transcript levels of pro-apoptotic YAP1 targets (**Figure 1H**), suggesting that the transcriptional increase in pro-apoptotic gene expression in response to the loss of CLN3 is conserved in a cell type prominently affected in Batten disease.

We next determined if the pro-apoptotic signaling resulting from absence of CLN3 was also observed *in vivo*. For that, we employed the CLN3^Δ7/8^ mice and their WT littermates, and analyzed the expression of YAP1 pro-apoptotic targets in two different brain regions, hippocampus and thalamus, which have been found to progressively deteriorate in Batten disease patients (Hochstein et al., 2022). Importantly, we observed an increase in the transcript levels of pro-apoptotic YAP1 target genes *Tp73*, *Dr5 and Cd68* were up-regulated in hippocampus (**Figure 1I**) and thalamus (**Figure 1J**) of 12-month old CLN3^Δ7/8^ mice compared to WT animals.

To verify that the pro-apoptotic signaling triggered by loss of CLN3 was indeed due to YAP1, we transiently silenced YAP1 by siRNA in HEK293 CLN3-KO cells (**Supplementary Figure S2G**), and observed a decrease in the transcript levels of YAP1 pro-apoptotic targets (**Figure 1K**), indicating that the pro-apoptotic signaling observed in CLN3-KO cells is downstream of YAP1. To determine if the increase in pro-apoptotic YAP1 transcriptional targets translates into increased apoptosis susceptibility, we assessed the cleavage of PARP, a substrate of caspase-3 and a well-known marker of apoptosis (Chaitanya et al., 2010), in CLN3-KO and WT cells. Under basal conditions, we observed similarly low levels of cleaved PARP in WT and CLN3-KO cells (**Figure 1L**), indicating low apoptotic activity in basal conditions. However, when the cells were subjected to a mild treatment with the apoptosis inducer staurosporine (2 µM, 4h), we noted a much higher increase of cleaved PARP in CLN3-KO cells than in WT cells (**Figure 1L**). These results suggest that, while CLN3-KO cells in basal conditions are still protected from apoptosis, these are more susceptible to cell death when subject to mild stress, likely due to YAP1 pro-apoptotic signaling.

### YAP1-dependent pro-apoptotic signaling in CLN3-KO is mediated by its interaction with p73

YAP1 activity is regulated by post-translational modifications, which determine its intracellular localization and transcriptional activity. In particular, YAP1 nuclear localization and pro-apoptotic activity have been associated with the phosphorylation of its tyrosine 357 (Y357) residue (Levy et al., 2008; Raghubir et al., 2021; Sugihara et al., 2018). Therefore, we tested whether phosphorylation of this residue (pYAP1^Y357^) was affected in CLN3-KO cells. We observed an increase in pYAP1^Y357^ in whole cell extracts of CLN3-KO cells (**Figure 2A**). Furthermore, we observed a robust enrichment of pYAP^Y357^ in the nucleus of CLN3-KO cells, while barely detected in the nucleus of WT cells (**Figure 2B**). To confirm that increased pYAP1^Y357^ is a specific consequence of CLN3-KO, we expressed CLN3-GFP in CLN3-KO cells, and observed a decrease in pYAP1^Y357^ (**Supplementary Figure 3A**). Furthermore, pYAP^Y357^ nuclear levels were also reduced in CLN3-KO cells expressing CLN3-GFP (**Supplementary Figure 3B**). To test if this result was also observed *in vivo*, we stained sections of CLN3^Δ7/8^ and WT mice brain with anti-pYAP^Y357^ antibody. Importantly, we found increased number of pYAP^Y357^-positive cells in the hippocampus and in the thalamus (**Figure 2C**), as well as an increase in total levels of pYAP^Y357^ in CLN3^Δ7/8^ brain tissue compared to WT (**Figure 2D**). These results underscore the importance of pYAP^Y357^ in the activation of YAP1 in both human CLN3-KO cells and rodent CLN3-KO brain.

**Figure 2.**
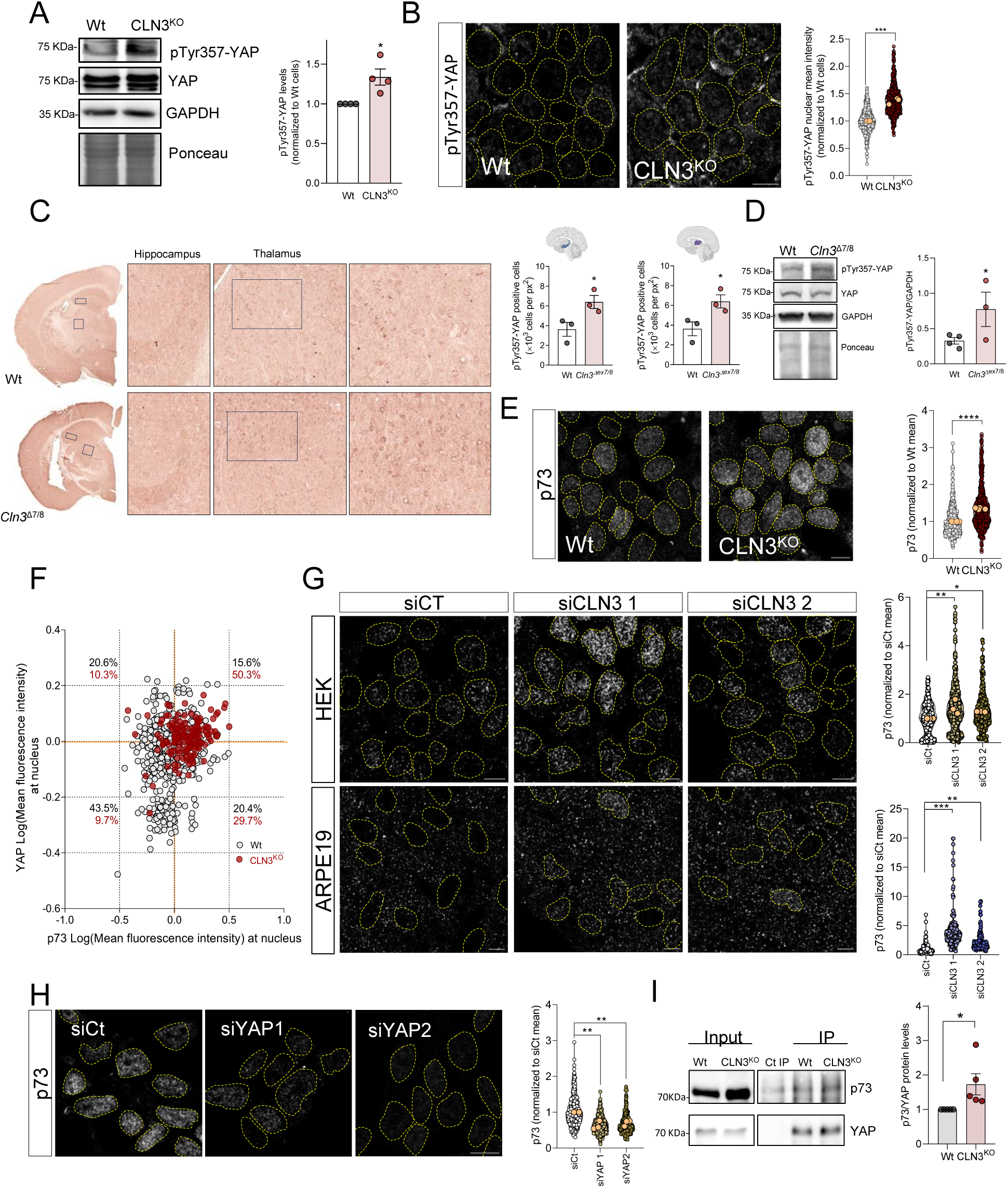
Loss of CLN3 promotes the interaction between YAP and p73. (**A**) Representative immunoblot image and quantification of YAP protein levels phosphorylated at the tyrosine residue 357 (pTyr357-YAP) in Wt and CLN3^KO^ cells. GAPDH and Ponceau were used as loading controls. (**B**) Confocal fluorescence images of Wt and CLN3^KO^ cells immunostained for pTyr357-YAP protein. Nucleus are outlined by the yellow dashed line using Hoechst staining. Scale bar 10µm. On the right, quantification of the mean intensity of YAP at the nucleus of at least 500 cells per each condition. The yellow dots represent the mean of each independent experiment. (**C**) Immunohistochemistry analysis of pTyr357-YAP staining of control (Wt) and *Cln3*^Δex7/8^ animals. On the right, quantification of the number of positive nucleus to pTyr357-YAP per the same area of hippocampus (first) and thalamus (second). (**D**) Representative immunoblot image and quantification of pTyr357-YAP in Wt and *Cln3*^Δex7/8^ animals. GAPDH and Ponceau were used as loading controls. (**E**) Confocal fluorescence images of Wt and CLN3^Kd^ cells immunostained for p73 protein. Nuclei are outlined by the yellow dashed lines using Hoechst staining. Scale bar 10µm. On the right, quantification of the mean intensity of p73 at the nucleus of at least 500 cells per each condition. (**F**) Correlation between the intensity levels of nuclear YAP (x axis) and p73 (y axis) proteins. At least 150 cells were analysed. (**G**) Confocal fluorescence images and quantification of HEK293T (upper panels and graph) and ARPE19 (bottom panels and graph) parental cell lines treated with scramble or CLN3-siRNA and immunostained for p73 protein. Nuclei are outlined by the yellow dashed line using Hoechst staining. Scale bar 10µm. All the results are mean±SEM of at least three independent experiments. (**H**) Confocal fluorescence images of CLN3^KO^ cells transfected with a control or two siRNA against YAP protein and immunostained for p73 protein. Nuclei are outlined by the yellow dashed lines using Hoechst staining. Scale bar 10µm. On the right, quantification of the mean intensity of p73 at the nucleus of at least 450 cells per each condition. (**I**) Representative immunoblot image and quantification of p73 co-immunoprecipitated with YAP protein in Wt and CLN3^Kd^ cells. The results are mean±SEM of four independent experiments. In the violin plots, yellow dots represent the mean of each individual experiment. Statistical analysis with one-way ANOVA followed by Dunnett’s multiple comparisons test (**G** and **H**), and Student’s *t*-tests when comparing two experimental groups. * p<0.05; ** p<0.01; ***p<0.001. * p<0.05; *** p<0.001; **** p<0.0001.

Curiously, the phosphorylation of Y357 is also reported to increase the interaction between YAP1 and p73, which is important for YAP1 pro-apoptotic signaling (Levy et al., 2008). We therefore tested if p73 levels were also increased in the nucleus of CLN3-KO cells. Our results show an increase in p73 nuclear localization in CLN3-KO cells (**Figure 2E**). To further explore if the nuclear localizations of YAP1 and p73 were correlated, we co-stained cells with antibodies against YAP1 and against p73, and observed that percentage of cells with increased YAP and p73 levels, increases from ∼15% in WT to >50% in CLN3-KOs (**Figure 2F**, upper-right quadrant). Accordingly, we transiently silenced CLN3 in HEK293 and ARPE19 cells, and observed an increase in p73 nuclear localization in both cell lines (**Figure 2G**), confirming that p73 nuclear localization responds specifically to loss of CLN3 rather than and adaptive mechanism in the CLN3-KO cells. Next, to determine if the accumulation of p73 in the nucleus was caused by YAP1, we transiently silenced YAP1 in CLN3-KO cells, and observed a robust decrease in the nuclear localization of p73 (**Figure 2H**). These results further underscores that there is an increase in both YAP1 and p73 in the nucleus in the absence of CLN3, the latter being dependent on YAP1.

Given that the p73 nuclear localization is YAP1-dependent, we tested if p73 nuclear localization may be mediated by protein-protein interaction with YAP1. We immunoprecipitated p73 from whole cell extracts of WT and CLN3-KO cells, and observed an increase in YAP1 associated with p73 in CLN3-KO cells (**Figure 2I**). Overall, these results show that the loss of CLN3 results in increased phosphorylation of YAP1 at Y357, which promotes its nuclear localization and interaction with p73.

Finally, given that loss of CLN3 affects the functioning of lysosomes (Nyame et al., 2024), we sought to clarify if the increase in pYAP^Y357^ and interaction with p73 may be a general consequence of lysosomal dysfunction, or a specific event caused by loss of CLN3. To this end, we tested two human cell lines with prominent lysosomal defects, namely HeLa cells deficient in alpha-glucosidase (GAA-kd; enzyme essential for glycogen degradation in lysosomes) or lysosomal proteases cathepsin B (CTSB-kd), which we characterized previously (Yambire et al, 2020; Agostini et al., 2024). We observed that these lines have lower levels of pYAP^Y357^ compared to the respective scrambled control cells (**Supplementary Figure S3C**). Furthermore, there was no enrichment of p73 in the nucleus of GAA-kd or CTSB-kd compared to the scrambled controls (**Supplementary Figure S3D**). Because lysosomal defects can also result in perturbations of the autophagy pathway, we tested if silencing Atg5, a protein involved in the early stages of autophagosome formation, would affect pYAP^Y357^ signaling. In Atg5-silenced HeLa cells, we observed no changes in the levels of pYAP^Y357^ (**Supplementary Figure S3E**). Altogether, these results suggest that the activation of YAP1 signaling is a specific consequence triggered by loss of CLN3 and not a generic consequence of impaired lysosomal function, or decreased autophagy.

### YAP1 activation in CLN3-KO cells depends on c-Abl activity

Given the importance of Y357 phosphorylation for the role of YAP1 in CLN3-KO cells, we sought to determine the mechanism leading to this phosphorylation. Considering that phosphorylation of YAP1 residue Y357 is mediated by the c-Abl tyrosine kinase (Levy et al., 2008), we started by assessing total c-Abl levels in whole cell extracts of WT and CLN3-KO cells, and observed an increase in CLN3-KO (**Figure 3A**). Because c-Abl continuously shuttles between the nucleus and the cytoplasm (Taagepera et al., 1998), we next determined if its nuclear localization is affected in CLN3-KO cells. Interestingly, we detected a robust increase in nuclear c-Abl in CLN3-KO cells when compared to the WT cells (**Figure 3B**). To confirm that c-Abl was responding specifically to the absence of CLN3, we expressed CLN3-GFP in the CLN3-KO cells, and observed a robust decrease in c-Abl nuclear localization (**Supplementary Figure S4**). We then sought to verify *in vivo* if c-Abl was also up-regulated in the absence of CLN3, and observed a prominent increase in c-Abl staining in both hippocampus and thalamus of CLN3^Δ7/8^ mice (**Figure 3C**).

**Figure 3.**
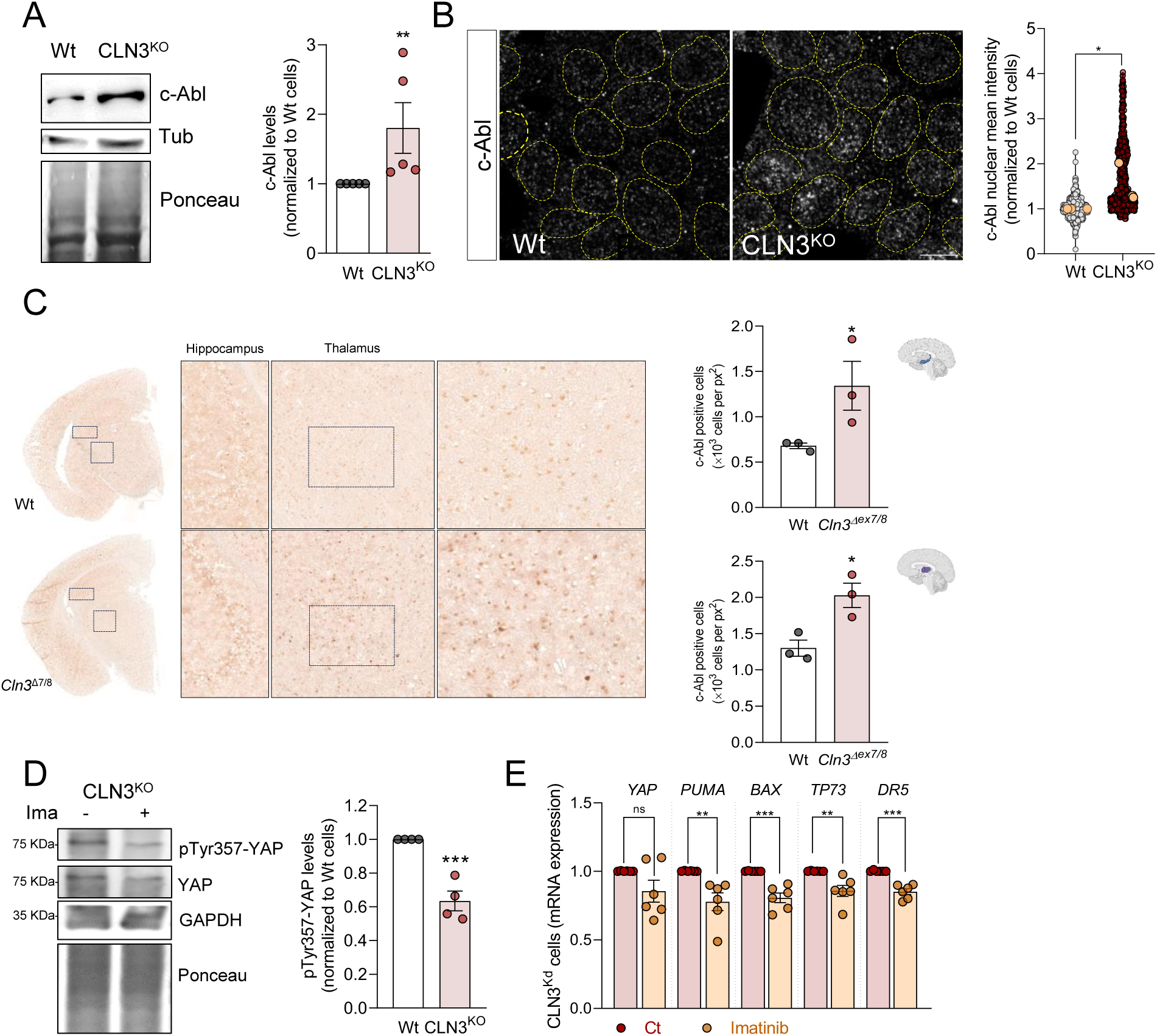
Loss of CLN3 triggers activation of c-Abl. (**A**) Representative immunoblot image and quantification of c-Abl protein levels in Wt and CLN3^KO^ cells. Tubulin and Ponceau were used as loading controls. (**B**) Confocal fluorescence images of HEK293T WT and CLN3^KO^ cells immunostained for c-Abl protein. Nucleus are outlined by the yellow dashed line using Hoechst staining. Scale bar 10µm. On the right, quantification of the mean intensity of c-Abl at the nucleus from at least 100 cells per condition in each independent experiment, with a total of at least 500 cells. The yellow dots represent the mean of each independent experiment. (**C**) Immunohistochemistry analysis of c-Abl staining of control (Wt) and *Cln3*^Δex7/8^ animals; Right panels are cropped images from thalamus; Quantification of the number of positive nucleus to c-Abl in the total area from the hippocampus (upper panel) and thalamus (bottom panel). (**D**) Representative immunoblot image and quantification of pTyr357-YAP in CLN3^KO^ cells control or treated for 24h with 0.1 µM imatinib (c-Abl inhibitor). GAPDH and Ponceau were used as loading controls. (**E**) mRNA levels of YAP target genes involved in apoptosis in HEK293T CLN3^KO^ cells untreated or treated for 24h with imatinib. All the results are mean±SEM of at least three independent experiments. * p<0.05; ** p<0.01; ***p<0.001. * p<0.05; *** p<0.001; using unpaired t-test.

Next, we tested if c-Abl was driving YAP1 hyperactivity by treating CLN3-KO cells with the c-Abl inhibitor imatinib (0.1μM, 24h). Our data revealed decreased levels of pYAP^Y357^ in whole cell extracts of CLN3-KO cells treated with imatinib (**Figure 3D**), without change in YAP1 total levels. Importantly, we also measured the transcript levels of YAP1 pro-apoptotic targets, and observed a decrease in imatinib-treated CLN3-KO cells (**Figure 3E**). These results show that the activity of YAP1 and its pro-apoptotic signaling in CLN3-KO cells are dependent on c-Abl activity.

### Activation of c-Abl in CLN3-KO cells in response to nuclear envelope instability

Considering that DNA damage is one of the major stimuli leading to c-Abl activation (Shaul & Ben-Yehoyada, 2005) and given that our transcriptome data indicated DNA damage/repair pathways being affected in CLN3-KO cells (**Figure 1B**), we next investigated nuclear homeostasis in response to CLN3-loss-of-function (**Figure 4A**). We first evaluated γH2Ax, a marker of unresolved DNA double-strand breaks, p53 and phospho-ATM^Ser1981^, both involved in DNA repair signaling, and observed a robust increase in the levels of these three proteins in CLN3-KO cells (**Figure 4B**). We also observed an increase in total levels of p53 and γH2Ax in the nucleus of CLN3-KO cells (**Figure 4C** and **Supplementary Figure S5A**). Moreover, by expressing CLN3-GFP in CLN3-KO cells, we were able to normalize the accumulation of double-strand breaks, assessed by quantification of γH2Ax (**Supplementary Figure S5B**). Furthermore, when we transiently silenced CLN3 in ARPE19 and HEK cells, we also observed an increase in both γH2Ax (**Supplementary Figures S5C-D**). This result confers further validity to the pathway analysis of the CLN3-KO transcriptome, which identified ATM signaling (**Figure 1B**), a key pathway of DNA damage response, as one of the most enriched pathways in the CLN3-KO transcriptome. Furthermore, we found a perturbation of the cell cycle in CLN3-KO cells, with increased retention of the cells in G1 phase (**Supplementary Figure S5E**), which is compatible with increased DNA damage and the cellular need to augment the time to repair DNA lesions. These findings then raise the question of what is the root of the DNA damage observed in the CLN3-KO cells.

**Figure 4.**
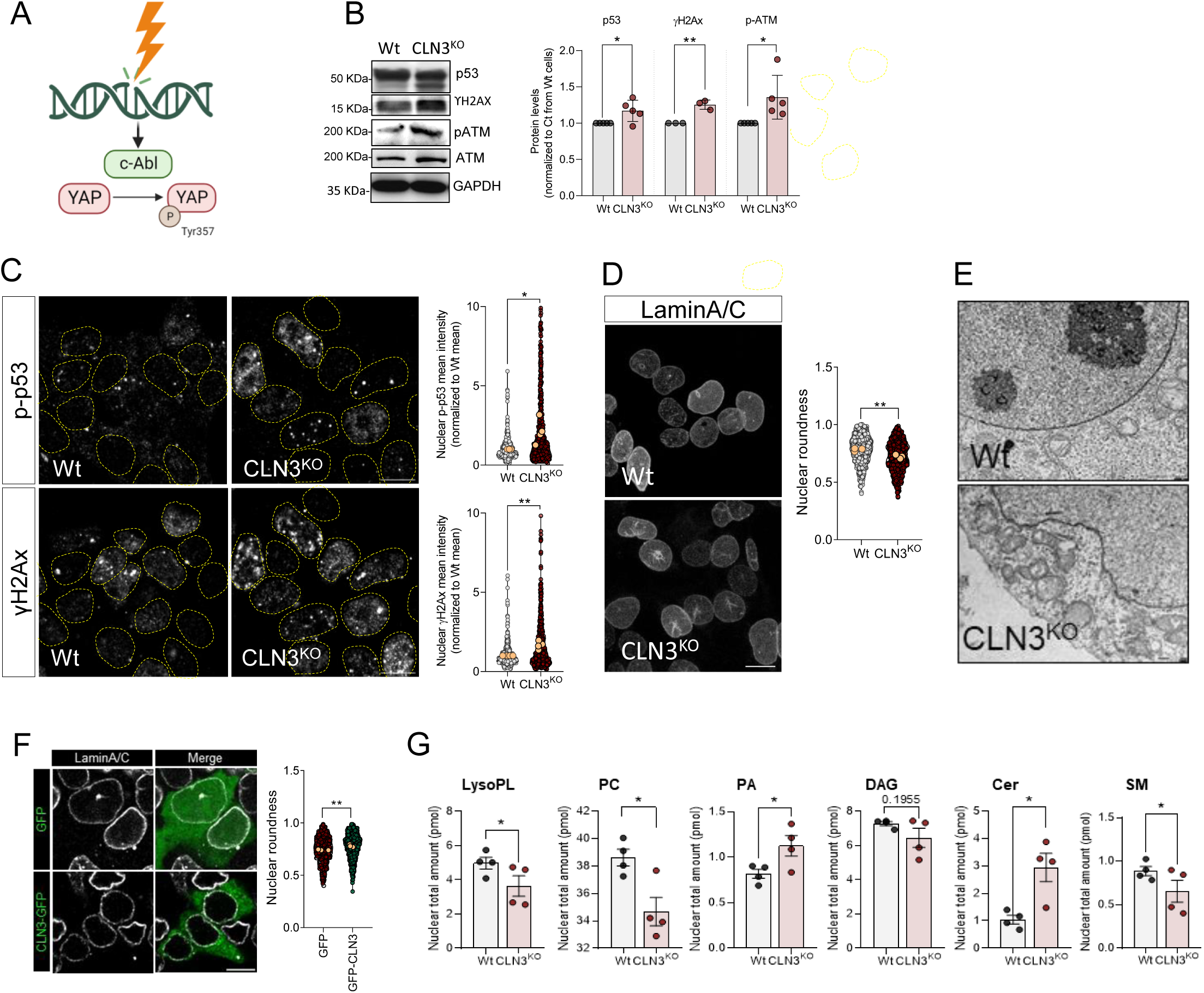
Accumulation of DNA damage in CLN3-KO cells drives c-Abl and nuclear dysmorphism. (**A**) Schematic representation of c-Abl activation in response to DNA damage. C-Abl directly phosphorylate YAP at the position Y357 in response to DNA damage. (**B**) Representative immunoblot image and quantification of p53, γH2AX, p-ATM and ATM protein levels in Wt and CLN3^KO^ cells. GAPDH was used as loading controls. (**C**) Confocal fluorescence images of HEK293T Wt and CLN3^KO^ cells immunostained for p-p53 and γH2AX proteins. Nucleus are outlined by the yellow dashed line using Hoechst staining. Scale bar 10µm. Graphs are the quantification of the mean intensity of p-p53 (upper panel) and γH2AX (bottom panel) at the nucleus from at least 100 cells per condition in each independent experiment, with a total of at least 450 cells. (**D**) Confocal fluorescence images of HEK293T Wt and CLN3^KO^ cells immunostained for LaminA/C proteins. Scale bar 10µm. Graphs represent the quantification of nuclear morphology parameters in HEK293T Wt and CLN3^KO^ cells immunostained for LaminA/C: circularity (up, right), aspect ratio (up, left), roundness (down, left) and solidity (down, right). (**E**) Transmission electron microscopy of HEK293T Wt and CLN3^KO^ cells. (**F**) Confocal fluorescence images of HEK293T CLN3^KO^ cells transfected with GFP or GFP-CLN3 and immunostained for LaminA/C proteins. Scale bar 10µm. Graphs represent the quantification of nuclear morphology parameters in HEK293T Wt and CLN3^KO^ cells immunostained for LaminA/C: circularity, aspect ratio, roundness and solidity. (**F**) Lipid composition of nuclear extracts isolated from HEK293T Wt or CLN3^KO^ cells. LysoPL= lysophospholipids, PC phosphatidylcholine, PA = phosphatidic acid, DAG= diacylglycerol, Cer= ceramide, SM=sphingomyelin. In the violin plots, yellow dots represent the mean of each individual experiment. All the results are mean±SEM of at least three independent experiments. * p<0.05; ** p<0.01; ***p<0.001. * p<0.05; *** p<0.001; using unpaired t-test.

During the course of our observation of many cellular phenotypes by imaging, we noticed that the nuclear shape of the CLN3-KO cells often seemed irregular (dysmorphic), while the WT cells seemed to mostly have ‘regular’ elliptic nuclei. To systematically address this question, we immunostained the WT and KO cells against Lamin A and Lamin C, two proteins found in the nuclear lamina. Interestingly, we observed that the number of dysmorphic nuclei was robustly increased in CLN3-KO cells, presenting a decrease in roundness (**Figure 5D**). To further explore this result, we performed electron microscopy on these cells, and again observed that while the nuclear envelope of the WT cells was mainly elliptic, there were many perturbations in the nuclear envelope of CLN3-KO cells (**Figure 5E**). Remarkably, nuclear morphology in CLN3-KO cells is normalized by expression of CLN3-GFP (**Figure 5F**).

**Figure 5.**
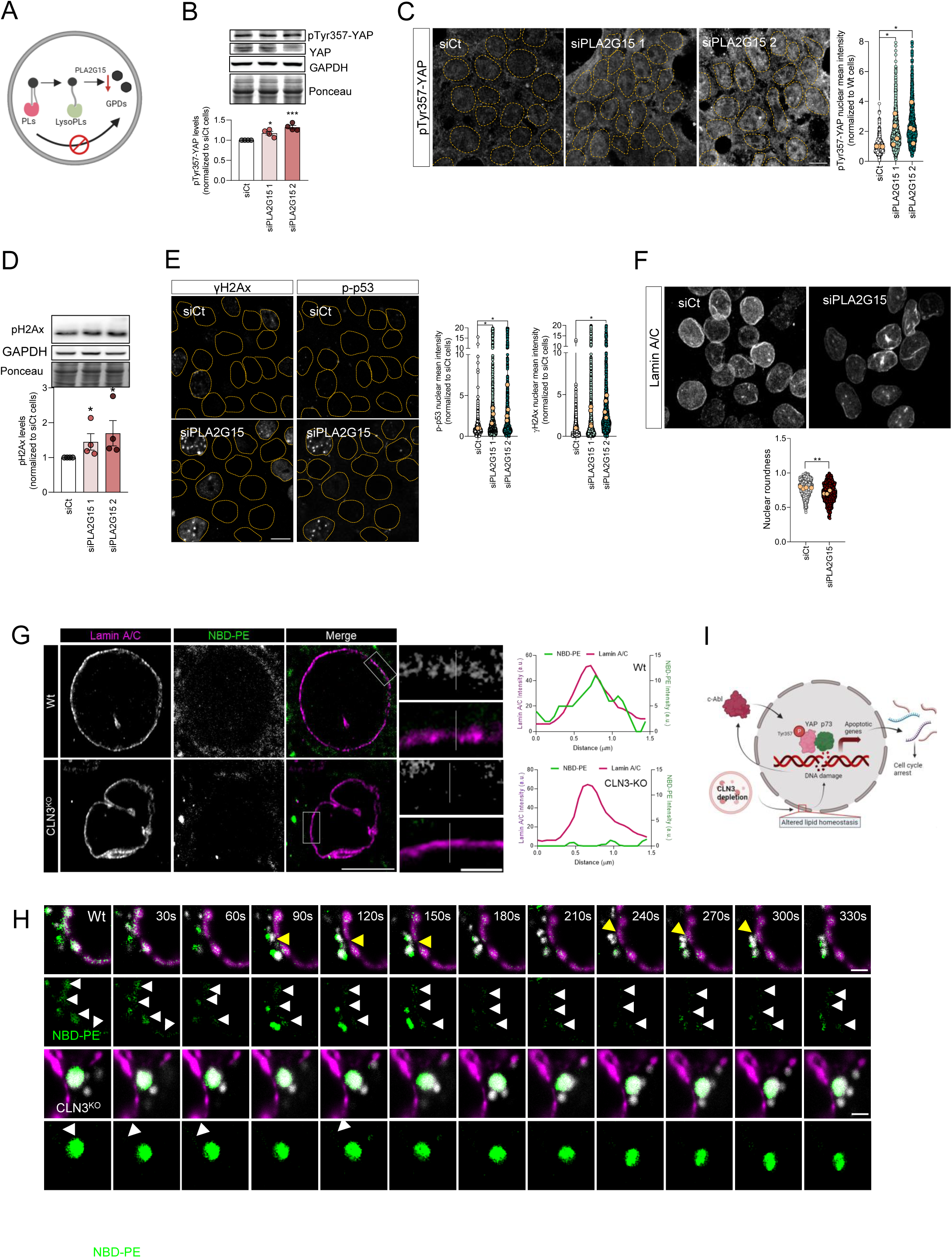
Inhibition of glycerophosphodiesters generation at the lysosome has the same effect on DNA damage and nuclear morphology as loss of CLN3. (**A**) Schematic representation of glycerophosphodiesters (GPDs) synthesis inhibition by affecting the activity of lysosomal lipases that mediate the hydrolysis of phospholipids into GDPs. (**B**) Representative immunoblot image and quantification of pTyr357-YAP protein levels in Wt cells silenced for YAP, using two different siRNA against lysosomal lipase A2 (PLA2G15). Ponceau was used as loading control. (**C**) Confocal fluorescence images and quantification of HEK293T Wt and cells depleted from PLA2G15 using two siRNA immunostained for pTyr357-YAP. Nucleus are outlined by the yellow dashed line using Hoechst staining. Scale bar 10µm. **(D)** Representative immunoblot image (**above**) and quantification (**below**) of γH2AX protein levels in Wt and CLN3^KO^ cells. Ponceau was used as loading controls. **(E)** Confocal fluorescence images and quantification (left for p-p53, right for γH2AX) of HEK293T parental cells transfected with siCT or siRNA against PLA2G15 using two different siRNA. Nucleus are outlined by the yellow dashed line using Hoechst staining. Scale bar 10µm. **(F)** Confocal fluorescence images of HEK293T Wt and CLN3^KO^ cells immunostained for Lamin-B proteins. Scale bar 10µm. The graphs represent the quantification of nuclear morphology parameters in HEK293T Wt and CLN3^KO^ cells immunostained for Lamin-A/C: circularity, aspect ratio, roundness and solidity. In the violin plots, yellow dots represent the mean of each individual experiment. **(G)** Representative confocal images of control and CLN3-depleted cells loaded for 24h with liposomes. To track the lipids distribution, cells were incubated with liposomes contain 1,2-myristoyl-sn-glycero-3-phosphoethanolamine-N-(7-nitro-2-1,3-benzoxadiazol-4-yl) (NBD-PE). At the upper panels, cells were stained for Lamin A/C. The graphs represent the fluorescence intensity profile of NBD-PE and Lamin A/C along the white line crossing the selected nuclear region (white line) in the inset. Scale bar 10µm and 2µm in the insets. **(H)** Schematic representation of our working model. **(I)** Timelapse images of live HEK293T Wt and CLN3^KO^ cells transfected with Lamin A-mRFP. For lysosomal labelling, cells were loaded with dextran AlexaFluor 647. Yellow arrows point near proximity of lysosomes and Lamin A, and white arrows highlight NBD-PE at nuclear envelope. Scale bar 2µm. All the results are mean±SEM of at least three independent experiments. * p<0.05; ** p<0.01; ***p<0.001. * p<0.05; *** p<0.001; using unpaired t-test.

We then investigated the reasons why the nuclear envelope might be perturbed in CLN3-KO cells. Recently, CLN3 was shown to be involved in the efflux of glycerophosphodiesters (GPDs) from the lysosome (Laqtom et al., 2022). GPDs are the product of lysosomal phospholipid degradation (Fowler and de Duve, 1969). We therefore hypothesized that the lysosomal accumulation of GPDs in conditions of CLN3 loss of function could impact the lipid composition of other organelles, namely the nucleus. To test this, we performed a lipidomics analysis in nuclear extracts from CLN3-KO and WT cells. Our data shows that lysophospholipids were significantly decreased in CLN3-KO nucleus, together with phosphatidylcholine and sphingomyelin. We also observed increased levels of phosphatidic acid and ceramides in the CLN3-KO nuclear extracts (**Figure 5G** and **Supplementary Figure S5F**). These results, in particular the decreased levels of phosphatidylcholine, lysophospholipids and sphingomyelin in CLN3-KO nuclei, suggest that the absence of CLN3 triggers changes in the composition of nuclear membranes that may result in genome instability and DNA lesions.

To further demonstrate that the lack of GPDs efflux from the lysosomes was causative of the nuclear phenotypes observed in CLN3-KO cells, we silenced the lysosomal phospholipase A (PLA2G15), which is the first enzyme processing glycerophospholipids in the lysosome (**Figure 5A**). In the absence of PLA2G15, no GPDs would be expected to be generated in the lysosomal lumen (**Supplementary Figure 6A**). By stopping the pathway upstream of CLN3, we can test if the decreased efflux of GPDs from the lysosomes is sufficient to perturb the nuclear shape, trigger DNA damage and, consequently, activation of YAP1 pro-apoptotic signaling. Interestingly, we found that silencing PLA2G15, which catalyzes the first step of lysosomal GPDs generation, was enough to increase the total levels of pYAP^Y357^ (**Figure 5B**) and in its nuclear localization (**Figure 5C**), as well as an increase in γH2Ax (**Figure 5D-E**). Of note, p53 was also enriched in the nucleus of PLA2G15-silenced HEK cells (**Figure 5E**). Moreover, by measuring nuclear morphology in PLA2G15-depleted cells, we also revealed an increase in nuclear dysmorphism in PLA2G15-silenced cells (**Figure 5F**), quantified as a decrease in roundness, similar to CLN3-KO cells.

Finally, we wanted to test if the GPDs released from the lysosomes could be found in the nuclear envelope. To this end, we added liposomes containing NBD-PE, with NBD linked to the ethanolamine group, to the WT and CLN3-KO cells (**Supplementary Figure S6B**). In agreement with previous findings, CLN3-KO cells presented increased NBD accumulation in lysosomes (**Figure 5G** and **Supplementary Figure S6C**). Notably, the nuclear envelope area, defined by lamina A/C immunostaining, presented NBD signal in WT cells, while no fluorescent signal was detected in the nuclear envelope area of CLN3-KO cells (**Figure 5G**). Interestingly, live imaging of lysosomal and NBD-PE dynamics at the nuclear rim (cytoplasmic regions in the proximity of the nuclear envelope) revealed an increase in lysosomal movement in control cells, while the lysosomes in CLN3-KO cells were enriched in NBD-PE and less mobile (**Figure 5G**, **Movie 1** and **Movie 2**). Notably, we also observed high level of ‘kiss-and-run’ events between lysosomes and nucleus, which can suggest an important role of CLN3 and lysosomal phospholipids in modulation of lysosome-nucleus contact sites (**Figure 5G-H**).

Altogether, these results show that the efflux of GPDs from the lysosome is necessary for DNA homeostasis and maintenance of the nuclear shape. When GPDs are not released from the lysosomes, as in PLA2G15-silenced cells or in CLN3-KO cells, then the nuclear envelope composition changes, nuclei became irregular and accumulate DNA damage, and promote c-Abl activation and YAP1-dependent pro-apoptotic signaling (**Figure 5I**).

## DISCUSSION

We show here that loss of CLN3, the lysosomal protein affected in Batten disease, leads to DNA damage and pro-apoptotic signaling in human cells and in mouse brain. CLN3 is involved in the lysosomal export of lysosomal phospholipid digestion products, specifically glycerophosphodiesters (GPDs; the “heads” of phospholipids after removal of the acyl chains) (Lagtom et al., 2022). Importantly, considering that lysosomal GPDs accumulation inhibits other enzymes of lysosomal phospholipid catabolism (Nyame et al., 2024), we silenced the first enzyme of the lysosomal phospholipid degradation pathway, PLA2G15. With this approach, we shut down the generation of degradation products of phospholipids, and therefore avoided that they accumulate and affect other lysosomal pathways. The silencing of PLA2G15 resulted in a perturbation of the nuclear envelope similar to CLN3-KO, with nuclear dysmorphism and DNA damage. The inability to release GPDs from the lysosome, either because of no export (lack of CLN3) or due to ablation of lysosomal phospholipid digestion (PLA2G15 silencing), results in a modification of the nuclear envelope lipidome, nuclear dysmorphism and impaired DNA homeostasis with accumulation of DNA damage.

Despite the recent finding that CLN3 protein is involved in the pathway that exports glycerophosphodiesters (GPDs) from the lysosomes (Laqtom et al., 2022), it is still unclear what are the consequences of the impairment of lysosomal GPDs export for cellular lipid homeostasis. Here, we show that the GPDs released from the lysosomes can be found on the nuclear envelope in healthy cells, but not in CLN3-KO cells. This result suggests that despite the GPDs can be soluble in the cytoplasm, those that are exported from the lysosomes may be rapidly metabolized in close proximity to the lysosome. Once exported from the lysosomes, GPDs are further metabolized by glycerophosphodiesterases (GDEs). There are seven GDEs in mammals (Corda et al., 2014), with different intracellular localizations, several of which are associated with cellular membranes in proximity of the nucleus. It is therefore plausible that GPDs released from lysosomes are metabolized in membranes near the lysosome upon export. We observe transfer of labelled GPDs from the lysosome to the nucleus in WT cells, but It remains an open question how do the products of lysosomal phospholipid degradation reach the nuclear envelope.

A membrane contact site between the nuclear envelope and the lysosome could facilitate the transfer of both GPDs and phospholipid degradation products from lysosomes to the nuclear envelope. The nucleus-lysosome contact sites have not been described in mammals, despite the vacuole-nucleus contacts have been described in yeast (Kvam & Goldfarb, 2007). We observed transient kiss-and-run contacts between the lysosomes and the nucleus, which might also be related to transfer of metabolites (for example, endosomes use kiss-and-run to transfer iron to mitochondria). It is also possible that the GPDs are transferred from the lysosomes to the endoplasmic reticulum (ER), via lysosome-ER contact sites, and then laterally diffuse towards the nuclear envelope, which is continuous with the ER. The involvement of lysosome-ER contact sites in the regulation of the lipid composition of the nuclear envelope and of DNA maintenance is supported by the presence of DNA damage in the subventricular zone of NPC1-KO mice (Seo et al., 2012). NPC1 is involved in lysosomal cholesterol and sphingomyelin efflux (Seo et al., 2012), and its loss-of-function results in decreased membrane contact sites between the lysosomes and the ER (Höglinger et al., 2019). Decreased association between lysosomes and the ER impacts the lipid composition of ER membranes (Agostini et al., 2024). Given that the nuclear envelope is a subdomain of the ER membranes, it is possible that changes in the lipid composition of the ER translate into modified lipid composition of the nuclear envelope. The pathway followed by GPDs or GPD-containing lipids exported from the lysosome towards the nuclear envelope warrants further investigation.

The lack of GPDs export from lysosomes in CLN3-KO cells would suggest a decreased availability of GPDs for salvage towards phospholipid synthesis. We observed that phosphatidylcholine, the most abundant phospholipid in the nuclear membrane, was present in lower concentrations in the nucleus of CLN3-KO cells. The decrease in sphingomyelin in CLN3-KO nucleus shows that other metabolites supplied by lysosomes are lacking in CLN3-KO nuclear envelope, suggesting that either the lysosomal accumulation of GPDs in CLN3-KO cells affects cholesterol and sphingomyelin export, or that the loss of transient kiss-and-run association between lysosomes and nucleus in CLN3-KO cells deprives the nucleus of multiple metabolites. A recent study of CLN3 proximity proteomics identified interactions between CLN3 and several subunits of the BORC complex (Calcagni’ et al., 2023),), which is involved in lysosomal positioning (Pu et al., 2015), raising the possibility that loss of CLN3 may affect lysosomal kiss-and-run association with the nucleus and GPDs export by independent synergistic pathways which culminate in modified lipid content of the nuclear envelope and nuclear dysmorphism.

Perturbations in the nuclear envelope homeostasis have been linked to premature aging disorders, chromosomal rearrangements and DNA damage, and are associated with pathologies such as laminopathies, cancer and cardiac disorders (Kalukula et al., 2022). The lipid composition of the nuclear envelope is poorly understood, and it remains unclear which mechanisms link the stability of DNA with the nuclear envelope composition. In the fission yeast *Schizosaccharomyces pombe*, it was shown that a ceramide synthase contributes to nuclear envelope integrity but the role of this process in DNA homeostasis was not explored (Hirano et al., 2023). However, *S. pombe* is a “closed mitosis” organism, because its nuclear envelope does not disassemble during mitosis, and therefore not a good model for mammalian nuclear envelope homeostasis. Furthermore, nuclear envelope tubules have been recently implicated in DNA repair (Shokrollahi et al., 2023), but it remains to be determined how the lipid composition of the nuclear envelope regulates this nuclear tubulation process.

The detection of nuclear dysmorphism and DNA damage markers downstream of impairment of lysosomal GPDs export opens the possibility for novel therapeutic strategies for patients with Batten disease caused by mutations in CLN3. Our data show that one of the proximal consequences of DNA damage is the activation of the kinase c-Abl, and that inhibition of c-Abl ablates YAP1 pro-apoptotic signaling. It remains to be tested *in vivo*, in models of CLN3-loss-of-function, if inhibition of c-Abl would delay the onset of symptoms or their progression. Strikingly, there are several commercially available drugs that target c-Abl and, if effective in CLN3-loss-of-function, they might constitute a therapeutic approach for Batten disease, which is currently incurable.

## MATERIALS AND METHODS

### Cell culture

Human embryonic kidney HEK293T cells were cultured in Dulbecco’s modified Eagle’s medium (DMEM, Thermo Fisher Scientific, Gibco, 12800082) supplemented with 10% FBS (Pan Biotech; P40-47500) and Penicillin/Streptomycin (100 U/ml:100 μg/ml), at 37°C under 5% CO2.

### Antibodies

The following antibodies were used: anti-YAP (Cell Signalling, 4912S; IB 1:1000); anti-pTyr357-YAP (Sigma, Y4645; IB 1:1000, IF 1:100); anti-p73 (Cell Signaling Technology, 14620S; IB 1:1000, IF 1:100); anti-cAbl (Cell Signalling 2662P; IB 1:1000, IF 1:100); anti-H2Ax (Cell Signalling, 9718S; IB 1:1000, IF 1:100); anti-p53 (Proteintech, 10442-1-AP; IB 1:1000, IF 1:100); anti-p-p53 (Cell Signalling, 9286P; 1:1000, IF 1:100); anti-mouse monoclonal anti-Tubulin (Sigma-Aldrich, T6199; IB 1:1000); anti-rabbit monoclonal anti-GAPDH (Cell signalling, 5174T; IB 1:1000); mouse polyclonal anti-hnRNPA2B1 [B-7] (Santa Cruz, sc-374053; IB:1000); anti-cleaved-PARP (Cell Signalling, 5625P, IB 1:1000); anti-ATM (Cell Signalling, 2873; IB 1:1000); anti-p-ATM (Ser1981) (D6H2) (Cell Signalling, 5883; IB 1:1000); anti-lamin A/C (Santa Cruz, 376248; IF 1:100); HRP-conjugated goat anti-mouse IgG (Cell Signalling, 7076S; IB 1:5000); HRP-conjugated goat anti-rabbit IgG (Cell Signalling, 7074P2; IB 1:5000); Alexa647-conjugated goat anti-rabbit IgG (Invitrogen, A21245; IF 1:500); Alexa568-conjugated goat anti-mouse IgG (Invitrogen, A11031; IF 1:500); mouse monoclonal anti-LAMP1 [H4A3] (development studies hybridoma bank, AB_2296838; IB 1:1000, IF 1:100); anti-CLN3 (kindly given by Alessia Calcagni’ (Calcagni’ et al., 2023); anti-LC3 (ab48394; IB 1:1000, IF 1:100); anti-PLA2G15 (Proteintech, 10863-2-AP; IB 1:1000).

### DNA constructs

CLN3-GFP plasmid was obtained from Addgene (#78110), a kind gift from Dr. Thomas Braulke.

### Generation of CLN3-KO HEK cells

To knock out CLN3 in HEK cells, we used a CRISPR–Cas9 approach. A guide RNA (gRNA) corresponding to the first exon of CLN3 protein was designed in CLC Work-bench 7 based on its genomic DNA sequence available on the Ensembl (Yates et al., 2020) (Forward 5’-CACCTGCGATGGGAGGCTGTGCAGGCTCG-3’ and Reverse 5’-AAACCGAGCCTGCACAGCCTCCCATCGCA-3’; IDT Technologies). Plasmid pSpCas9(BB)-2A-Puro (PX459) was digested with BbsI. The band at 9kb, corresponding to the total size of the plasmid (9175 bp), was cut out from the gel and purified utilizing NucleoSpin Gel and PCR Clean-up kit, and subsequently ligated with the gRNA, followed by transformation of *E. coli* strain XL10-Gold ultracompetent cells, from which the plasmid was subsequently purified. To generate CLN3-KO in HEK cells, transfection of parental HEK cells was performed using Lipofectamine 3000 according to the supplier’s instructions. After 48 hours the media in transfected and natural puromycin resistance control plates were changed to a growth medium with 2μg/ml of puromycin added to start antibiotic selection. Selection media were changed every 2-3 days until all Wt cells in the control plate were dead as determined by observation under the microscope. Single-cell colonies were obtained by flow cytometry at the Core Facility Cell-Sorting of the University Medical Center Göttingen, and DNA was sequenced to determine colonies with CLN3-KO.

### Mice

Cln3Δ7/8 knock-in mice generation was previously described (Cotman et al., 2002). Mice were genotyped using the following primers:

5’-TTTGTTCTGCTGGGAGCTTT-3’, 5’-CAGTCTCTGCCTCGTTTTCC3’-,

5’-GACAAGAGCACTGAGGAAGAT-3’, 5’-AGGAAGGAATGGGGAGACTGA-3’, resulting in a 600bp band in the case of Cln3Δ7/8 mice, and a 450bp band for the wild type allele. Mice used for experiments were maintained in a C57BL/6J strain background.

All experiments were approved by the Committee on Animal Care at Baylor College of Medicine (protocol AN-5280) and conform to the legal mandates and federal guidelines for the care and maintenance of laboratory animals.

### Cell treatments and liposomes

After 24 hours of seeding, cells were treated with: 2µM staurosporine for 4 hours; 0.1μM imatinib for 24h; and liposomes 1.5 mM of total lipids for 24h.

Liposomes were prepared as previously described (Martins et al., 2019). Briefly, a lipid film of DPPC:PE-NBD (9:1) was performed followed by hydration with PBS. The resulting multilamellar vesicles were extruded using a 100 nm pore size filters (Nucleopore, Whatman, Springfield Hill, U.K.) using a Extrusor from Lipex [Biomembranes](https://www.sciencedirect.com/unsupported_browser) (Vancouver, British Columbia, Canada).

### Cell transfection and siRNA-mediated knockdown

DNA and siRNA transfections were performed using Lipofectamine 3000 according to the manufacturer’s instructions. siRNA target sequences were CLN3 (Dharmacon, siGENOME SMARTpool M-004233-02-0005), YAP1 (Thermo Fisher Scientific, s94586) and PLA2G15 (IDT, hs.Ri.PLA2G15.13.1, hs.Ri.PLA2G15.13.2, hs.Ri.PLA2G15.13.3). After 24h of cell seeding, 20 nmol/L siRNA was complexed with the transfection reagent and incubated at 37°C. After an overnight incubation, transfection media was replaced with fresh media and the experiments performed after 72h. Non-targeting control sequences (Ambion) were used as controls.

### Immunofluorescence and cell imaging

Cell lines were grown on gelatine-coated coverslips were fixed in 4% (w/v) PFA for 10 min, followed by 20 min permeabilization and blocking with PBS with 0.05% (w/v) saponin and 1% (w/v) BSA (blocking solution). Coverslips were then incubated overnight with primary antibodies prepared in the blocking solution at 4°C. Next, coverslips were washed with PBS and incubated with Alexa Fluor-conjugated secondary antibodies for 1 h, at room temperature. Nuclei were stained with Hoechst. Finally, coverslips were mounted with Mowiol 4-88. In order to confirm their specificity, all the antibodies were tested through immunoblot analysis and secondary antibody controls, where primary antibodies were not added. Images were acquired on a confocal microscope (Zeiss LSM 710; Carl Zeiss AG) by using a Plan-Apochromat 63x/1.4 oil DIC M27 objective. For each experimental condition, at least 5 different fields were imaged. The imaging of immunostained cell was performed using the same settings.

For live imaging, cells were seeded in Ibidi chamber. After an overnight transfection, lipid and Dextran Alexa Fluor 647 loading, cells were washout for 4 hours with complete DMEM without phenol red supplemented with 10mM HEPES. Live cells were imaged using the LSM710 confocal system and a 63x/1.4NA objective. A single stack was acquired by time-lapse imaging, through the acquisition of one z-stack image per 30 seconds for a total of 10 min.

### Immunohistochemistry

After embedding brains in OCT, blocks were sectioned coronally at 10 μm. For immunohistochemistry procedures, sections were washed with 1x Tris-buffered saline (TBS1X), blocked with 5% goat serum, 0.25% Triton in TBS for 60 minutes, then treated for 10 minutes with the Bloxall solution to quench endogenouse peroxidases (BLOXALL, vectorlab,SP-6000). Next, sections were washed in TBS and incubated overnight at 4°C with primary antibodies diluted in the blocking solution (5% goat serum, 0.25% Triton in TBS). The next day sections were washed in TBS, and incubated with the MACH4 HRP-Polymer (M4U534 G, Biocare Medical) for 1H. Slides were rinsed in TBS 1X and immune reactions were revealed using the Peroxidase (HRP) Substrate, NovaRED (SK-4800, Vector Labs). Sections were washed and mounted with permanent aqueous mounting medium (Biorad, BUF058B). Anti-Abl, Sigma SAB4500045 (1:200), Anti pYap, Sigma Y4645 (1:100)

### Image analysis

Fluorescent image analysis was performed on the original data using Fiji Image J1 software (version 1.53 q). For quantification of nuclear levels of YAP, pTyr357-YAP, p73, c-Abl, p-p53, nuclear regions were defined by applying a threshold to Hoechst staining and using the ‘analyze particles’ plug in from Fiji (definition of our region of interest, ROIs). The background imaged was measured and removed from each nucleus. For nuclear shape assessment, z-stack images were acquired from Lamin A/C immunostaining and the resulting z-project image was analyzed.

### RNAseq

RNA sequencing was performed as described (Murdoch et al., 2016a). RNAseq data analysis was performed using Partek Software Suite. RNAseq data was aligned to the reference genome mm10 by the BoWtie algorithm, and the transcripts were quantified using the human GRCh37 Ensembl assembly as reference. For differential expression analysis, Bonferroni multi-test correction was applied, and adjusted p-value<0.05 was considered significant. Ingenuity Pathway Analysis was used for assessment of transcription factor activity, as described (Murdoch et al., 2016b; Yambire, Fernandez-mosquera, et al., 2019; Agostini et al., 2024). The data is deposited at Gene Expression Omnibus (GSE265831).

### Nuclear extracts

Nuclear extracts were performed using the Nuclear/cytosol fractionation kit (Abcam, AB289882) following the manufactures indications.

### Lipidomics

MS-based lipid analysis was performed at Lipotype GmbH (Dresden, Germany).

### Quantitative RT-PCR

RNA was extracted with NucleoBond kit isolation kit. Reverse transcription was performed using iScript cDNA synthesis kit (Bio-Rad, 170-8891). Quantitative PCR was performed in a 384-well plate (BioRad) using the SYBR green master mix (BioLabs, M3003E) using CFX Opus 384 Real-Time PCR System with One software (v2.2.2; Applied Biosystems). Hprt and ActA were used as housekeeping genes to normalize the expression. Target gene expression was determined by relative quantification (ΔΔCt method) to the housekeeping reference gene and the control sample. The primer sequences are indicated in Table S2.

### Immunoprecipitation

Cell lysates were prepared in RIPA buffer, supplemented with protease and phosphatase inhibitors (ThermoScientific, 78440). Immunoprecipitations were performed using 500μg of total protein lysates. The YAP1 primary antibody (0.5μg) was incubated with cell at 4°C under rotation overnight. Protein G Sepharose beads were then added and incubated for 1h, at 4°C. After washing, complexes were eluted in Laemmli buffer, boiled for 5 minutes at 95°C and analyzed by Western blot.

### Western blot

Cell extracts (50 µg of protein) were prepared using Laemmli buffer. Before the gel loading, samples were sonicated and boiled for 5 min at 95°C. Subsequently, samples underwent separation via sodium dodecyl sulfate polyacrylamide gel electrophoresis (SDS-PAGE) and were transferred onto nitrocellulose membranes. Ponceau staining was conducted for total protein evaluation, followed by washing. Membranes were washed and then blocked with 5% (w/v) non-fat milk in Tris-buffered saline-Tween 20 (TBST; 20 mM Tris, 150 mM NaCl, 0.2% (v/v) Tween 20, pH 7.6) during at least 30 min. Primary antibodies were incubated overnight at 4°C, succeeded by incubation with HRP-conjugated secondary antibodies for 1 h at room temperature. Protein detection was accomplished via chemiluminescence (ECL prime western blotting detection reagent, Cytiva, Amersham, RPN2236) using an ImageQuant 800 from Cytiva. Quantification was performed on unsaturated images using Image Lab software (Bio-Rad).

### Transmission electronic microscopy

HEK293T or HEK293T CLN3-KO cells were fixed with 2.5% glutaraldehyde in 0.1M sodium cacodylate buffer (Agar Scientific, pH 7.2) supplemented with 1mM calcium chloride. Sequential post-fixation was performed using 1% osmium tetroxide (Sigma), for 1.5 h, and contrast enhanced with 1% aqueous uranyl acetate, for 1h. After rinsing in distilled water, samples were embedded in 2% molten agar and then dehydrated in a graded ethanol series (50–100%), impregnated and embedded using an Epoxy embedding kit (Fluka Analytical). Ultrathin sections (70nm) were mounted on copper grids (300 mesh) and stained with lead citrate 0.2%, for 7min. All observations were carried out using a FEI-Tecnai G2 Spirit Bio Twin at 100 kV.

### Statistical analysis

All the experiments were repeated at least three times, using independent experimental samples and statistical tests as specified in the figure legends. The independent replicates were obtained from different passages of the same cell line. Statistical analysis was conducted using Graphpad Prism version 8.0.1. All distributed data are displayed as means ± standard error of the mean (SEM). A p-value lower than 0.05 was considered statistically significant, and significance levels were indicated as indicated with *p < 0.05, **p < 0.01, ***p < 0.001, ****p < 0.0001.

## ACKNOWLEDGEMENTS

We thank Dr. Nicolas Lemus for his expert assistance with RNAseq data normalization and processing, and the CNC microscopy facility. This work was supported by NIH R56AG082790-01 (to NR), FCT 2022.09311.PTDC, FCT ERC-Portugal, NCL-Stiftung; Welcome Trust Investigator Award in Science 224361/Z/21/Z, La Caixa, John Black Foundation (to I.M.). The intitial stage of this work was supported by ERC Starting Grant 337327 (to N.R.) and DFG SFB1190-P02 (to N.R. and I.M.). This project also received funding from the European Union’s Horizon 2020 research and innovation programme under grant agreement MIA-Portugal No 857524 and the Comissão de Coordenação e Desenvolvimento Regional do Centro - CCDRC through the Centro2020 Programme. The content is solely the responsibility of the authors and does not necessarily represent the official views of the National Institutes of Health.

## AUTHOR CONTRIBUTIONS

ND, AC, JP, SRF, NJH, TH, KW performed experiments. ND, AC, IM and NR designed experiments. ND, AC, SRF, JP, NJH, TH, TFO, HG, IM, AB and NR analyzed and interpreted data. NR designed the project. ND and NR prepared the figures and wrote the manuscript, which all authors commented on.

## Notes

### Competing Interest Statement

The authors have declared no competing interest.

### Summary of Updates

Correction of one typo on the name of one of the authors; correction of the institutional affiliations for three authors. Added one chapter that was missing in the methodology.

