## Supplementary Figures for "Loss of the lysosomal protein CLN3 modifies the lipid content of the nuclear envelope leading to DNA damage and activation of YAP1 pro-apoptotic signaling"

### Supplementary information

#### Cell cycle analysis by flow cytometry

Cell cycle from HEK293T control and CLN3-KO cells were synchronize through an overnight starvation using HBSS. After the synchronization, HBSS was replaced by complete medium for 0 and 9 hours, and then washed and fixed with cold ethanol (70%) for 15 min on ice. Cells were next washed and treated with ribonuclease. Propidium iodide (PI) was added with a final concentration of 30 µg/mL.

Data were acquired in a BD FACSCalibur flow cytometer (Becton Dickinson): PI fluorescence was measured in the PI channel (585/42 BP). The software for acquisition was BD FACSDiva, and data from at least 5000 cells were analysed with FlowJo.

#### Lysosomal pH and degradative activity

For lysosome pH, cells were loaded with dextran conjugated with FITC (100 µg/mL, Merck, 60842-46-8) and dextran conjugated with Alexa Fluor 647 (60 µg/mL, ThermoFisher, D22914) overnight, followed by a 4h chase before treatment with 100 nM bafilomycin for 1h. Before cells imaging, cells were treated for 15 min with Magic-red (MR, Bio-Rad, ICT937) at 37°C. Cells were live-imaged by confocal microscopy in presence of 1 mM of HEPES in complete DMEM without phenol-red.

To measure the fluorescence intensity ratio between FITC/ Alexa Fluor 647-dextran and MR/ Alexa Fluor 647, a threshold was applied to Alexa Fluor 647-dextran channel to define the lysosomes through the analyse particle macro from ImageJ Fiji software. The background was removed from the obtained fluorescence values.

### Figure S1 – Characterization of CLN3KO cell line

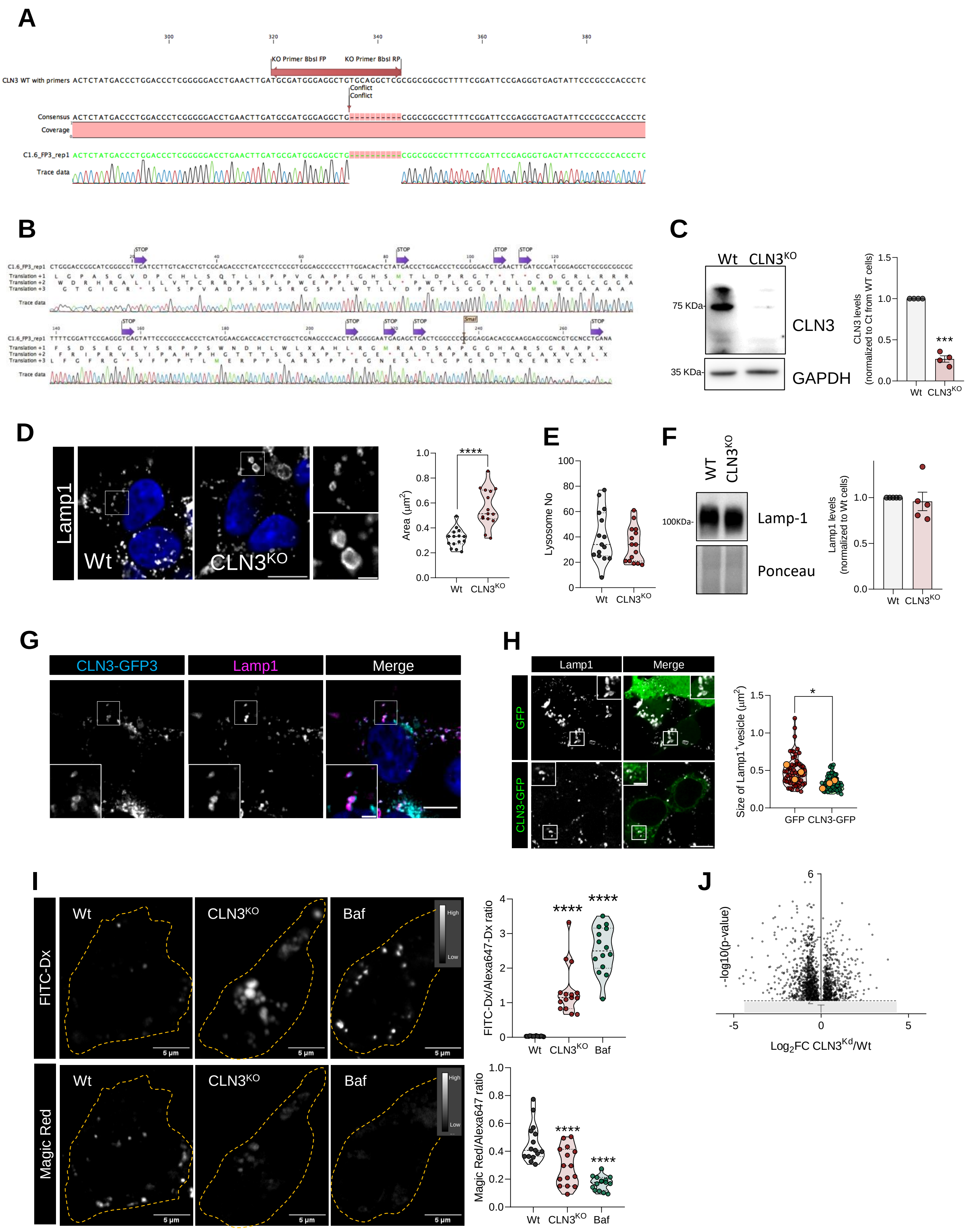

**Figure S1** Characterization of HEK293T CLN3-KO cell line.

**(A)** Schematic representation of sequence deleted in CLN3 encoding gene. **(B)** Putative translation sequences showing the premature stop codons. **(C)** Representative immunoblot image and quantification of CLN3 protein levels in Wt and $CLN3^{KO}$ cells. GAPDH and ponceau were used as loading controls. **(D)** Confocal fluorescence images of HEK293T Wt and $CLN3^{KO}$ cells immunostained for Lamp1 protein and the respective quantification of lysosomal area. Scale bar 10µm and 2µm in the inset images. **(E)** Quantification of total number per cell in HEK293T Wt and $CLN3^{KO}$ cells. **(F)** Representative immunoblot image and quantification of Lamp1 protein levels in Wt and $CLN3^{KO}$ cells. Ponceau were used as loading controls. **(G)** Confocal fluorescence images of cells HEK293T parental line transfected with CLN3-GFP and immunostained for Lamp1. Scale bar 10µm and 2µm in the inset images. **(H)** Confocal fluorescence images of cells HEK293T $CLN3^{KO}$ transfected with CLN3-GFP and immunostained for Lamp1. Quantification of lysosomal size from at least 50 HEK293T $CLN3^{KO}$ cells transfected with GFP or CLN3-GFP. **(I)** Representative fluorescence images of Wt and $CLN3^{KO}$ cells loaded with FITC-dextran (Dx), sensitive to pH, and magic red fluorescence, a substrate of cathepsin B. Scale bar 10µm. The graphs represent the quantification of FITC-dextran (on the top) and magic red fluorescence (on the bottom) normalized to Alexa647-dextran, not pH sensitive. At least 15 cells were analysed. All the results are mean±SEM of at least three independent experiments. * $p<0.05$; ***$p<0.001$; **** $p<0.0001$; using unpaired t-test or Dunnett's multiple comparisons test. **(J)** Differentially expressed genes (DEG) between HEK293T wild type (Wt) and CLN3 knockdown ($CLN3^{KO}$) are represented as a volcano plot: log p-value adjTtest in the y-axis and log2FC in the x-axis. The transcripts overexpressed in KO are to the right of the plot, and on the left, the repressed genes.

### Figure S2

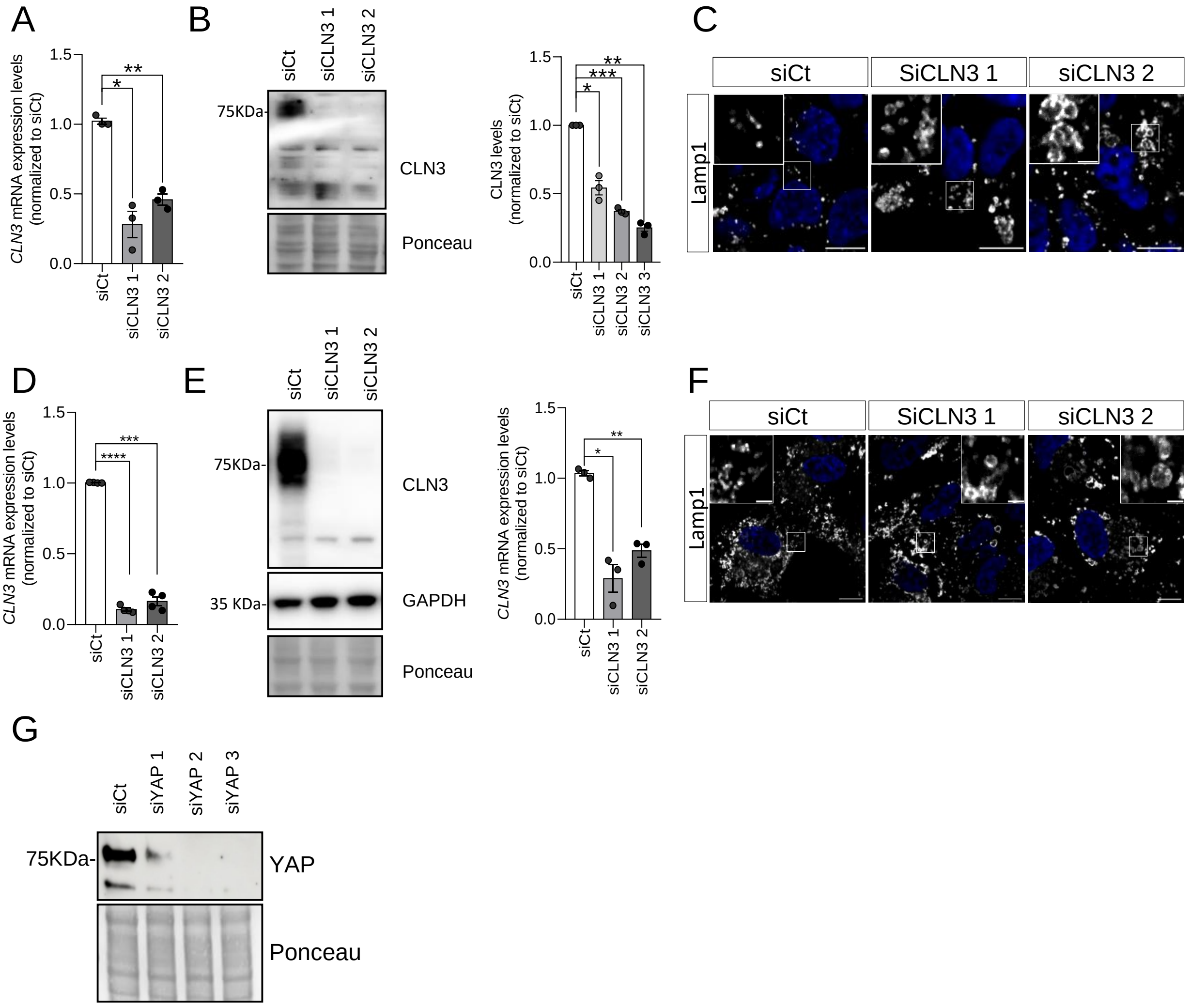

**Figure S2** Characterization of transient modulation of CLN3 and YAP levels in HEK293T parental line cell line.

(**A**) mRNA levels of CLN3 gene in HEK293T parental line transfected with siCt or two different siRNA against CLN3 transcript. (**B**) Representative immunoblot image and quantification of CLN3 protein levels in the conditions described in (**A**). Ponceau was used as loading controls. (**C**) Confocal fluorescence images of cells under the conditions described in (**A**). Scale bar 10µm and 2µm in the inset images. (**D**) mRNA levels of CLN3 gene in ARPE19 parental line transfected with siCt or two different siRNA against CLN3 transcript. (**E**) Representative immunoblot image and quantification of CLN3 protein levels in the conditions described in (**D**). Ponceau was used as loading controls. (**F**) Confocal fluorescence images of cells under the conditions described in **D**. Scale bar 10µm and 2µm in the inset images. (**G**) Representative immunoblot image YAP protein levels from HEK293T parental line transfected with siCt or three different siRNA against YAP transcript. Ponceau was used as loading controls. All the results are mean±SEM of at least three independent experiments. * $p<0.05$; ** $p<0.01$; *** $p<0.001$; **** $p<0.0001$; using Dunnett's multiple comparisons test.

### Figure S3

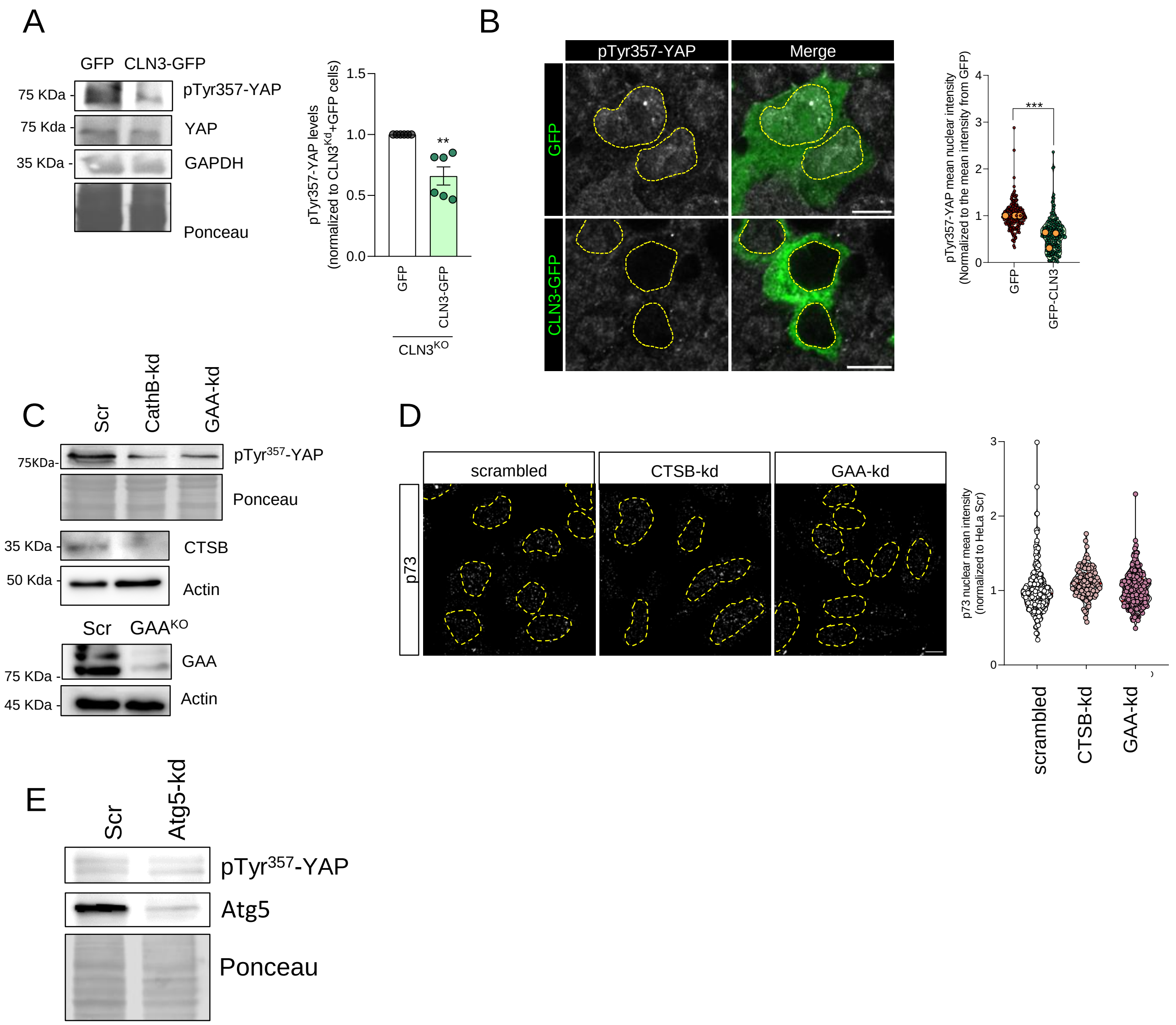

**Figure S3** Restoring CLN3 levels in HEK293T CLN3-KO cells led to a decrease in YAP phosphorylation at the Tyr357 residue.

(**A**) Representative immunoblot image and quantification of pTyr357-YAP in CLN3$^{KO}$ cells transfected with GFP or CLN3-GFP treated. GAPDH and Ponceau were used as loading controls. (**B**) Confocal fluorescence images and quantification of cells under the conditions described in **A** immunostained for pTyr357-YAP. Scale bar 10µm. At least 150 nucleus were analysed. The results are mean±SEM of at least three independent experiments. **p<0.01; ***p<0.001 using t-test. (**C**) Representative immunoblot image of pTyr357-YAP in control, Cathepsin-B knock-down (CTSB-kd) and acid alpha-glucosidase knock-down (GAA-kd) HeLa cells. Protein depletion was also confirmed by CTSB and GAA immunoblotting. Actin, GAPDH or Ponceau were used as loading controls. (**D**) Confocal fluorescence images and quantification of cells under the conditions described in **C** immunostained for p73. (**E**) Representative immunoblot image of pTyr357-YAP in control and Atg5 knock-down (Atg5-kd) HeLa cells.

**Figure S4**

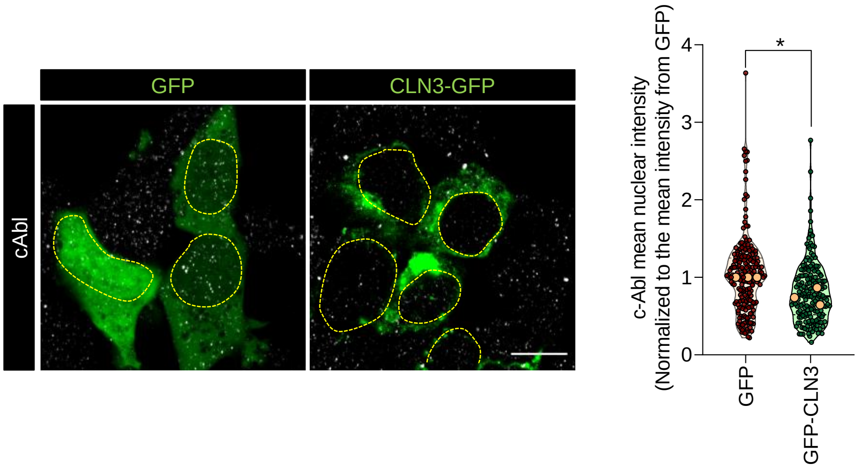

**Figure S4** Restoring CLN3 levels in HEK293T CLN3-KO cells led to a decrease in c-Abl recruitment to the nucleus.

(**A**) Confocal fluorescence images and quantification of $CLN3^{KO}$ cells transfected with GFP or CLN3-GFP treated and immunostained for c-Abl. Scale bar 10µm. At least 150 nucleus were analysed. All the results are mean±SEM of at least three independent experiments. * $p<0.05$; ** $p<0.01$; using t-test.

### Figure S5

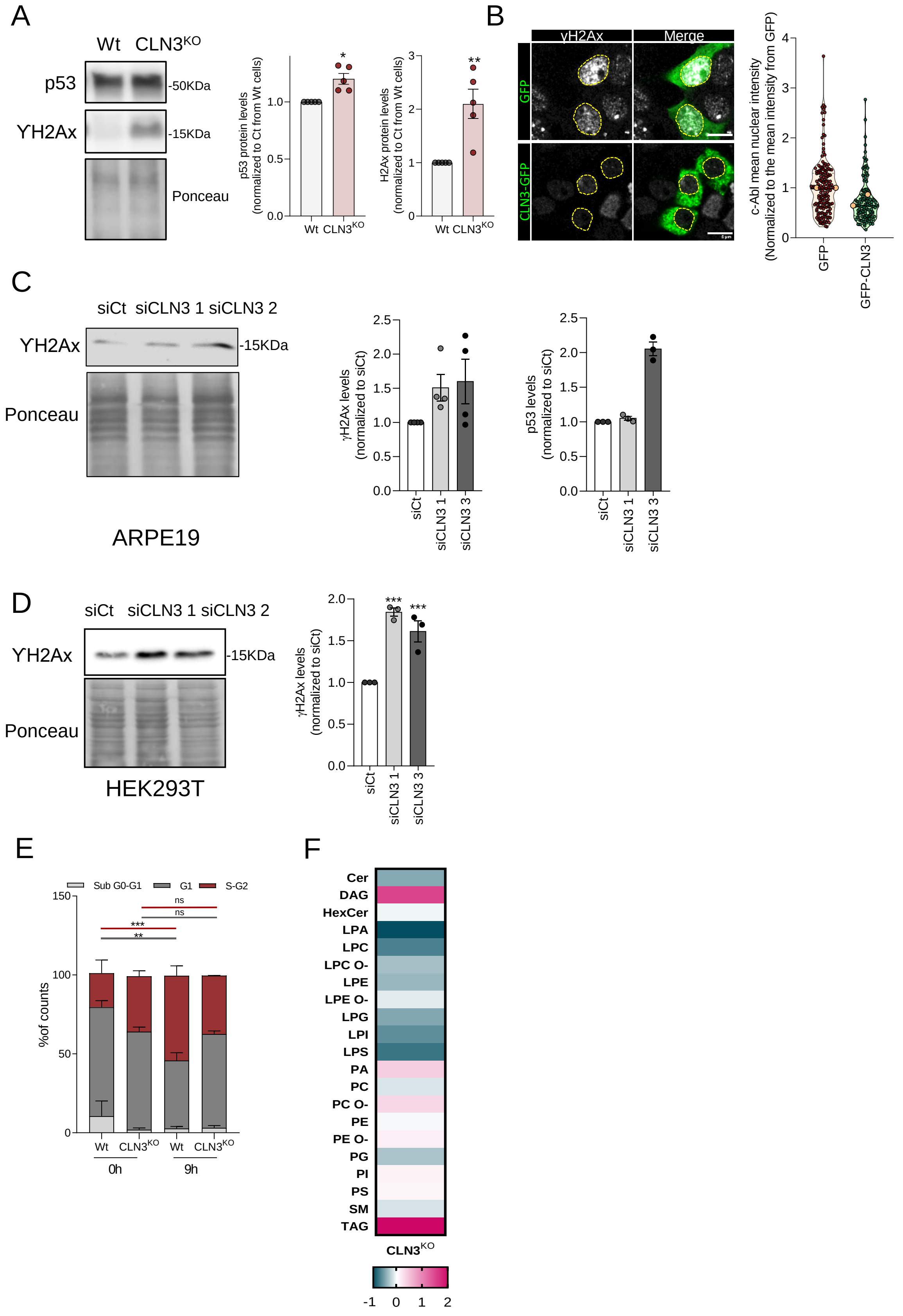

A
Wt CLN3KO
p53
-50KDa
YH2Ax
-15KDa
Ponceau
p53 protein levels (normalized to Ct from Wt cells)
H2Ax protein levels (normalized to Ct from Wt cells)
Wt CLN3KO
B
γH2Ax
Merge
GFP
CLN3-GFP
c-Abl mean nuclear intensity (Normalized to the mean intensity from GFP)
GFP
GFP-CLN3
C
siCt siCLN3 1 siCLN3 2
YH2Ax
-15KDa
Ponceau
ARPE19
γH2Ax levels (normalized to siCt)
p53 levels (normalized to siCt)
siCt
siCLN3 1
siCLN3 3
D
siCt siCLN3 1 siCLN3 2
YH2Ax
-15KDa
Ponceau
HEK293T
γH2Ax levels (normalized to siCt)
siCt
siCLN3 1
siCLN3 3
E
Sub G0-G1
G1
S-G2
%of counts
ns
ns
***
**
Wt CLN3KO Wt CLN3KO
0h
9h
F
Cer
DAG
HexCer
LPA
LPC
LPC O-
LPE
LPE O-
LPG
LPI
LPS
PA
PC
PC O-
PE
PE O-
PG
PI
PS
SM
TAG
CLN3KO
-1 0 1 2

**Figure S5** Loss of CLN3 function induces increase in DNA damage and cell cicle arrest possible by affecting nuclear lipidome.

(**A**) Representative immunoblot image and quantification of p53 and ϒH2AX in nuclear extracts from HEK293T parental and CLN3-KO cell line. Ponceau was used as loading controls. (**B**) Confocal fluorescence images and quantification of $CLN3^{KO}$ cells transfected with GFP or CLN3-GFP treated and immunostained for ϒH2AX. Scale bar 10µm. At least 150 nucleus were analysed. (**C-D**) Representative immunoblot image and quantification of p53 and ϒH2AX in ARPE19 (**C**) or HEK293T (**D**) parental cell line transfected with siCt or two siRNA against CLN3 transcripts. Ponceau was used as loading controls. (**E**) Cell cycle quantification of Wt and $CLN3^{KO}$ cells stained with propidium iodide (PI) after cells synchronization with an overnight starvation. The percentage of cells in each cycle phase was quantified by flow cytometry (three independent experiments). (**F**) Heat map of lipid classes affected in the nucleus of CLN3-KO when compared with HEK293T parental cell line. All the results are mean±SEM of at least three independent experiments. * $p<0.05$; ** $p<0.01$; *** $p<0.001$; using t-test or Dunnett's multiple comparisons test.

### Figure S6

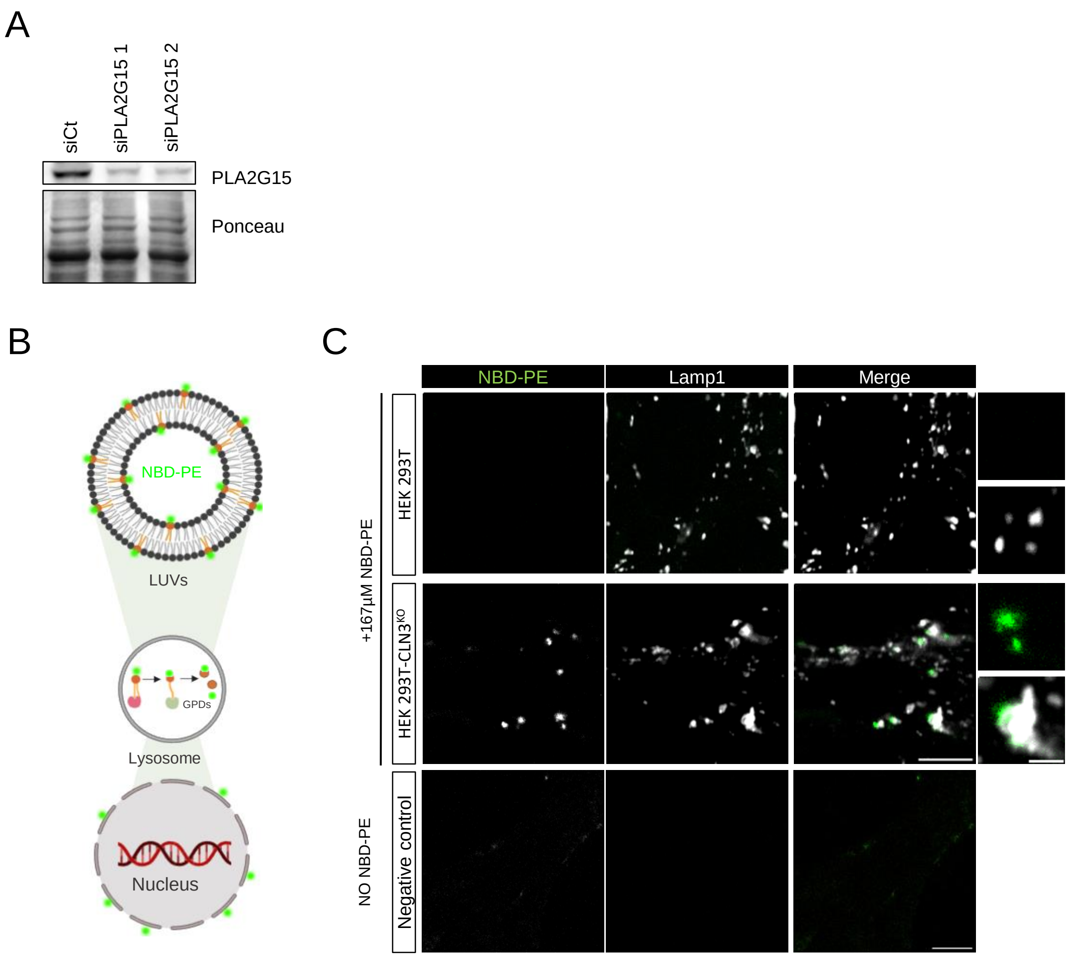

**Figure S6** Confirmation of PLA2G15 depletion and NBD-PE distribution in CLN3-KO cells.

(**A**) Representative immunoblot image of PLA2G15 in HEK293T parental cell line transfected with siCt or two siRNA against PLA2G15 transcripts. Ponceau was used as loading controls. (**B**) Schematic representation of the experimental approach using large unilamellar vesicles (LUVs) containing NBD-PE fluorescent lipids. LUVs will be internalized by endocytosis and lipids degradated by lysosomal lipases. We hypothesised that products resulting from fluorescent lipids degradation will be detected at nuclear envelope. (**C**) Representative confocal images of Wt and CLN3-KO cells loaded with NBD-PE-containing liposomes and immunostained for Lamp1. Scale bar 10µm and 2 µm in the insets.

**Table 1 – List of human primers and siRNA sequences.**

| *Primers* | *Sequences (5'-3')* |
|---|---|
| **h_PUMA_F** | ATCAATCCCATTGCATAGGTTTAG |
| **h_PUMA_R** | ACTAAGGCTGGGGCGCTTC |
| **h_TP53AIP1_F** | GGCTCAGACACACACACCT |
| **h_TP53AIP1_R** | GGCCTGTCTCTAAGCACTGT |
| **h_Bax_F** | ATGTTTTTCTGACGGCAACTTC |
| **h_Bax_R** | ATCAGTTCCGGCAACCTTG |
| **h_TP73_F** | CCCACCACTTTGAGGTCACT |
| **h_TP73_R** | GGCGATCTGGCAGTAGAGTT |
| **h_DR5_F** | GTGATTCAGGTGAAGTGGAGC |
| **h_DR5_R** | CGACCTTGACCATCCCTCTG |
| **hYAP1_F** | TAGCCCTGCGTAGCCAGTTA |
| **hYAP1_R** | TCATGCTTAGTCCACTGTCTGT |
| *siRNA* | *Sequences (5'-3')* |
| **h_siCLN3_1** | UUGUUCUUUCAAGGUCUAUUCUUUAUAGACCUUGAA |
| **h_siCLN3_2** | GCAGUACCGAUGGUACCAUAGCAUCUGGUACCAUCG |
| **h_siYAP_1** | ACGGUAGAUAUUACUGACAAUUCAUCAGAUAAUAU |
| **h_siYAP_2** | GCUGCCACCAAGCUAGAUUUUCUUUAUCUAGCUUGG |
| **h_siPLA2G15_1** | GUAUCUGGAUUCUGGCAAACUUUUAUUGCCAGAAUC |
| **h_SiPLA2G15_2** | AGACCGAAAGCUACUUCACAGAUUGUGAAGUAGCUU |

**Table 1 – List of murine primers.**

| | |
|---|---|
| S16-F | AGGAGCGATTTGCTGGTGTGG |
| S16-R | GCTACCAGGGCCTTTGAGATG |
| Cycloph-F | GGCAAATGCTGGACCAAACACAA |
| Cycloph-R | GTAAAATGCCCGCAAGTCAAAAG |
| mYAP1-RT-F | CGGCAGTCCTCCTTTGAGAT |
| mYAP1-RT-R | GGTCCTGCCATGTTGTTGTC |
| mTP53-RT-F | AACTATGGCTTCCACCTGGG |
| mTP53-RT-R | TGAGGGGAGGAGAGTACGTG |
| mBBC3-RT-F | GGATGGCGGACGACCTCAA |
| mBBC3-RT-R | TCGGTGTCGATGCTGCTCTT |
| mBAX-RT-F | ATCCAAGACCAGGGTGGCTG |
| mBAX-RT-R | TCACTGTCTGCCATGTGGGG |
| mTrp73-RT-F | TCACCTTCCAGCAGTCGAGC |
| mTrp73-RT-R | TGGATGGGGCATGTCTTAGCA |
| mDr5-RT-F | AGCCCATCAAGAGGACCCTG |
| mDr5-RT-R | AGGCTTGCAGTTCCCTTCTGA |
| mcd68-RT-F | TCAGCTGCCTGACAAGGGAC |
| mcd68-RT-R | GCAGCAAGAGGGACTGGTCA |
| mCln3ACFOR | CCTCCAGGAAAGGTGGACAG |
| mCln3ACREV | GCTCGAAAAGTCCCTGGTTG |
